# Riemannian Metric Learning for Alignment of Spatial Multiomics

**DOI:** 10.64898/2025.12.09.693237

**Authors:** Peter Halmos, Yufan Xia, Benjamin J. Raphael

## Abstract

Recent spatial technologies measure the transcriptome, epigenome, proteome, metabolome and other modalities from thousands of cells across a tissue. Most assays typically profile only one modality from a tissue slice, raising the question of how to align spatial data from heterogeneous feature spaces. While multiple approaches have been developed for multi-modal integration of single-cell datasets, few existing techniques perform spatial alignment across arbitrary modalities incorporating both spatial and feature information. We introduce Manifold Gromov-Wasserstein (`MGW`), a metric-learning framework that exploits the product structure of spatial multiomics to infer modality-specific Riemannian *pull-back* metrics with neural fields. `MGW` aligns Riemannian distances induced by these metrics via Gromov-Wasserstein optimal transport, yielding a hyperparameter-free cost across arbitrary modalities sharing a spatial base. The formulation enjoys theoretical invariances – including orthogonal transformations of the spatial and feature domains as well as global feature scalings. We demonstrate the advantages of `MGW` on multiple alignment tasks, including Stereo-Seq spatiotemporal transcriptomics of mouse embryo, Xenium and Visium spatial transcriptomics of colorectal cancer, and spatial metabolomics-transcriptomics from human striatum and kidney cancer. `MGW` recovers biologically meaningful correspondences and spatially coherent tissue structures, outperforming existing OT- and non-OT-based multi-modal baselines.

**Code availability:** Software is available at https://github.com/raphael-group/MGW

## 1 Introduction

Spatially-resolved transcriptomic (SRT) technologies (48; 42; 36) are a seminal advance towards a comprehensive understanding of spatial biology (2; 24), but they measure only a single of many biological modalities which define the state of a cell. Beyond transcription, a cell state is defined by modalities such as its metabolome, epigenome, and proteome. More recent spatial multiomics technologies combine measurements of the transcriptome with profiling of the epigenome (60) in spatial ATAC-Seq and spatial CUT & Tag-RNA-Seq, profiling of spatial metabolomics and histology (56), profiling of open chromatin and T-cell receptor sequences (43), and profiling of protein markers (4). However, most technologies only measure a single modality for a given slice. While modern computational techniques have offered great promise for understanding biology from a functional lens – from understanding cell-differentiation dynamics (46; 47; 45; 16) to spatial heterogeneity (51; 21; 13; 58; 6) and cellular niches (19; 14) – extending such questions to the spatial multiomic universe requires *alignment* across differing modalities.

Multiple works have tackled the problem of aligning spatial transcriptomics data (26; 7; 30; 59; 17). Techniques for spatial alignment have proven highly effective, yet universally face the difficult problem of how to weight spatial similarity against feature similarity. Some methods such as `STAlign` (7) avoid the question by relying on spatial information alone. Another popular approach called *fused Gromov-Wasserstein* (FGW) optimal transport (54), introduces a trade-off hyperparameter *α* between spatial and feature information. FGW was first introduced to spatial transcriptomics by `PASTE` (59) and adapted by many following works to account for partial alignment (30), unbalanced and semi-relaxed alignment (26; 17), and learning cell-fate landscapes (23). However, its hyperparameter is often difficult to tune and the framework is not extendable to multiple modalities.

Alignment of single-cell datasets from different modalities is also well-studied. Multimodal alignment presents a unique challenge, as these spaces may have differing dimensions, incomparable scales and geometries, and no common basis of coordinates on which points can be compared. However, aligning across heterogenous spaces is the most general and powerful form of alignment: it subsumes any notion of mode, from experimental batch to time point to technology. Works such as `SCOT` (12; 11), `SCOTv2` (10), and `moscot` (26) align single-cell datasets across modalities using Gromov-Wasserstein optimal transport, matching the modality-spaces based on their pairwise distance structures. In contrast, numerous non-OT approaches – based on variational autoencoders, nearest-neighbor anchoring, or matrix-factorization (18; 49; 57; 29) – perform multiomic integration by learning a shared latent representation, but typically do not return explicit cross-dataset alignments. A universal limitation of existing OT-based multimodal alignment methods is that they do not leverage spatial information and do not *learn* a cost function. Instead, they rely on Euclidean or graph distances computed symmetrically within each modality on the raw features.

One approach to spatial multi-modal integration is to learn joint embeddings of spatial multiomics via graph representations or autoencoders, as in `MultiGate` (34) and `SpatialMeta` (53) but these methods presume *pre-aligned* cells. Even when an alignment module is included for spatial multiomics (e.g. using `STAlign` (7) in (53)), registration operates on *coordinates*, not on the feature-space geometry. Another recent and parallel line of work has investigated learning of the *Riemannian metric* of the data-manifold (1; 44), with applications to learning the latent manifold with autoencoders (50; 38). However, no existing techniques have bridged learning modality-specific metrics for the underlying spatial-molecular manifold with alignment across heterogeneous spaces.

### Contributions

We introduce *Manifold Gromov-Wasserstein* (`MGW`), an algorithm to align spatial multiomics datasets across arbitrary modalities. The key insight of `MGW` is to learn a cost whose respective mapping across modality spaces preserves the *intrinsic manifold structure* of each space. `MGW` is based on the observation that a spatial ‘omics dataset can be viewed as a function from physical (Euclidean) space *E* ⊂ ℝ^2^ or ℝ^3^ to a high-dimensional feature, or modality, space ℳ. As each modality shares the same base space *E*, `MGW` utilizes a standard approach in Riemannian geometry for studying different manifolds sharing a common base space: the *Riemannian pull-back metric*, which pulls the geometry of the modality spaces back onto the common Euclidean space. To compute the pull-back metric, we represent each spatial-omic dataset with a *neural field*: an implicit map from physical space into the respective modality space. Finally, `MGW` computes an alignment using Gromov-Wasserstein Optimal Transport on the Riemannian distances in each modality. Manifold Gromov-Wasserstein offers a geometrically-unified technique for spatial multimodal alignment and is free of hyperparameters for weighting between spaces.

We show that `MGW` is a reasonable measure of “distance” between spatial-feature manifolds, with invariance to rigid-body transformations of the spatial coordinates and invariance to orthogonal transformations, offsets, and homogeneous scalings of the features. Additionally, `MGW` recovers GW on spatial coordinates when the fields are identity maps. We demonstrate that `MGW` outperforms existing spatial alignment methods across four different spatial-omics alignment tasks spanning five measurement technologies. `MGW` bridges neural representation and feature-learning with optimal transport, offering a natural way to *learn* costs for optimal transport to align across different modalities.

## 2 Methods

A spatial-omics dataset 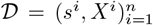 is a set of pairs of spatial locations *s*^*i*^ and associated features *X*^*i*^. For spatial multiomics alignment one has a common Euclidean base space *E* (e.g. *E* = ℝ^2^, ℝ^3^), and a pair of spaces ℳ ⊂ ℝ^*d*^, ⊂ 𝒩 ℝ^*p*^ representing the space for each measured modality (e.g. ℳ transcription, and 𝒩 metabolomics). We begin by introducing the general problem of aligning such multimodal data with a common Euclidean base space.

### Problem 1(Multimodal Alignment with Common Base Space.).

*Given spatial datasets* 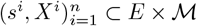 *and* 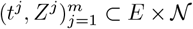 *with modality spaces* ℳ *and* 𝒩 *over a common Euclidean base E, and given a cross-space cost C*_*M*×*N*_ : (*E* × ℳ) × (*E* × 𝒩) → ℝ,^1^ *find a coupling* ***π*** *between the modalities* ^2^, *which minimizes the expected cost*

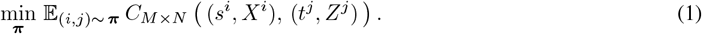

For fixed cost *C*_*M*×*N*_ (·, ·), Problem 1 is well-defined and the best alignment can be found using optimal transport (OT) (See (10) and (11)). The true challenge, then, is in determining or learning the most appropriate notion of cost (39). As different modality spaces exhibit incomparable dimensions, scalings, and geometries, identifying such a cost is difficult. A commonly used cost in spatial transcriptomics alignment is the Fused Gromov-Wasserstein (FGW) cost, which is defined to be a convex combination of a cost on feature-distances evaluated *across* datasets and a cost on spatial distances evaluated *within* a dataset (54; 59; 30; 26). This, however, poses existential limitations for Problem 1: FGW requires distances across the two modalities while its underlying cost functions are within one space, e.g. *ℓ*_2_ distances.

The key principle behind `MGW` is to not pre-define a cost, but to instead *learn* a cost which preserves the *intrinsic manifold structure* of each modality space. This is not only to map across heterogeneous spaces, but also to preserve non-linear manifold geometry. Even when two datasets exist in spaces which are not directly comparable, the two modalities may exhibit a common geometry when viewed relatively within each space. To define such a cost, we make two key assumptions on our data:

**(A1)** The modality spaces are *Riemannian manifolds* (ℳ, *g*) and (𝒩, *h*) equipped with associated Riemannian metrics to measure distances.

**(A2)** A pair of smooth mappings exist from the base Euclidean space into the respective modality spaces. We denote these by *ϕ* : *E* → ℳ and *ψ* : *E* → 𝒩.

We note (A1) automatically holds for the base Euclidean space (*E, g*^*E*^) with metric being the identity *g*^*E*^ = **id**. Given (A1) and (A2), the core question of our work then becomes the following:

> *Can we learn a cost between arbitrary modalities measured over Euclidean space which can preserve the intrinsic geometry of the underlying data manifolds?*

### 2.1 Manifold-Pullback and Pullback Metric

In this section, we describe how `MGW` learns modality-specific notions of distance and pulls them back to the underlying Euclidean space. To make this precise, we first recall a few preliminaries from Riemannian geometry. Given a smooth manifold *M*, a *Riemannian metric* or *metric tensor* is assigns to each point *p* ∈ *M* a symmetric, positive-definite bilinear form *g*_*p*_ : *T*_*p*_*M* × *T*_*p*_*M* → ℝ on the tangent space *T*_*p*_*M* such that the coefficients *g*_*ij*_(*p*) vary smoothly in every local coordinate chart. The metric acts as an inner product on tangent vectors in *T*_*p*_*M* and defines a local notion of length and angle. When *M* = ℝ^*d*^, the canonical choice *g*(*X, Y*) := *X, Y* yields the standard Euclidean geometry. The pair (*M, g*) is called a Riemannian manifold (see, e.g., (27; 28) for standard references).

To extend these objects to spatial-multiomics, we need to learn the geometry of the features over physical space. We note that a spatial dataset 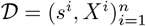 can be viewed as *n* samples 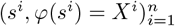 of tissue coordinates and a function *ϕ* : *E* → ℳ mapping the tissue coordinates (Euclidean space) into the modality space ℳ. We represent this function for each modality using an *implicit neural field* or *implicit neural representation ϕ* : *E* → ℳ, *ψ* : *E* → 𝒩. Fields offer a different perspective on spatial multiomics: rather than representing the data as a finite grid or matrix of spatial and feature coordinates (Figure 1), one views the data *implicitly* as a continuous field of features over physical space. Spatial multiomics data are well-suited to field-based parametrizations: gene expression or other molecular modalities vary smoothly and continuously across tissue (6), and exhibit non-linear *isodepth* coordinates along which expression changes with a differentiable gradient (6; 31). For `MGW`, the key value of the field-based representation is in capturing the spatial differential structure underlying modality variation over space. With access to spatial derivatives 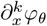, one can access essential geometric objects – gradients, curvature, and induced metric tensors.

**Figure 1:**
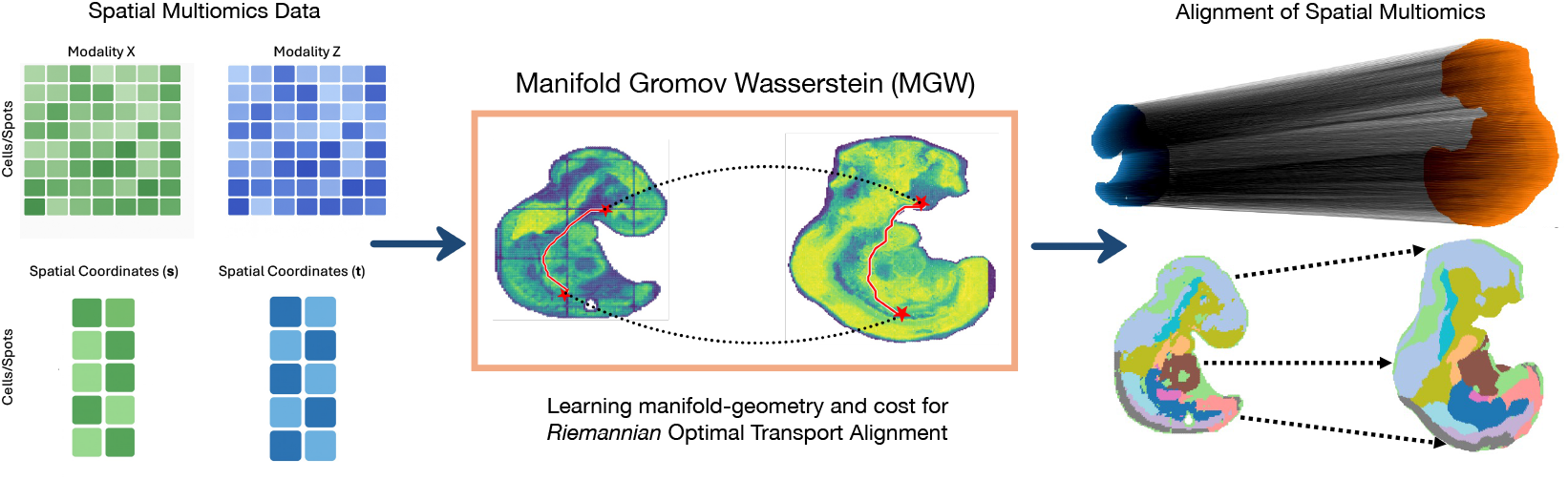
Overview of Manifold Gromov-Wasserstein (`MGW`). (Left) Two spatial multiomic datasets are input including both features for each modality and corresponding spatial coordinates. (Middle) `MGW` learns the intrinsic Riemannian geometry of each dataset with neural fields and aligns Riemannian distances. (Right) The alignment offers a non-linear mapping between cells measured in arbitrary modalities while preserving the intrinsic geometry of each dataset.

Following (6), we learn these field representations by minimizing the mean-squared loss

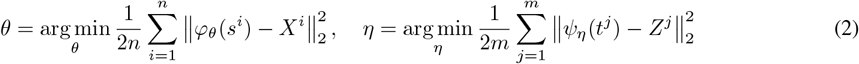

And taking *ψ* := *ψ*_*η*_, *ϕ* := *ϕ*_*θ*_ to be the implicit representations of each omics slice defined as a field defined over Euclidean space. For our architecture, we restrict to *C*^∞^ activations; in particular, we use the SiLU function (41; 35). This ensures that *ψ*_*η*_, *ϕ*_*θ*_ have smooth Jacobians and represent smooth submanifolds, when viewed as local parametrizations mapping from Euclidean space *E* into the respective modalities. Many Jacobians in neural representations of omics data play key roles: gene-over-gene Jacobians characterize temporal gene-regulatory networks (47) and gene-over-space or modality-over-space Jacobians (Definition 1) characterize the gradients and curvature of a modality over physical space (6; 31). As the modality-over-space Jacobian is central to this work, we recall it below.

#### Definition 1(Space-Modality Jacobian.).

*Let ϕ* : *E* → *M denote a field mapping spatial coordinates s* 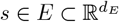 *to a feature-space M* ⊂ ℝ^*d*^. *At each point s, the* space-modality *Jacobian is the differential*

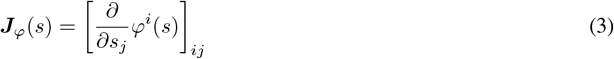

(3) encodes how spatial displacements change modality (feature) abundances. As *s* varies over *E*, ***J***_*ϕ*_() : *E* → *L*(*TE, T*_*ϕ*_*M*) defines a smooth *Jacobian tensor field* of order (1, 1) on *E*. In other words, each point *s* in *E* maps to a Jacobian encoding the manner in which spatial displacements change features at that point. Now, given the Jacobian (3), we define the *pullback metric* of the fields *ϕ, ψ* onto the Euclidean base *E*.

#### Definition 2(Modality Pullback Metric).

*Given a field ϕ with Jacobian field* ***J***_*ϕ*_(·) : *E* →*L*(*TE, T*_*ϕ*_*M*), *at a point x the Pullback-Metric induced by ϕ for modality M is*

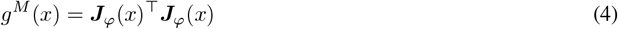

*A symmetric, positive-definite* (0, 2) *tensor on the tangent space T*_*x*_*E. The associated inner product is given by*

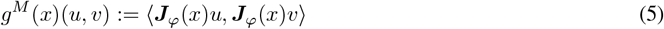

*Observe that g*^*M*^ (·) : *E* → *S*^2^(*T*^∗^*E*) *defines a smooth* (0, 2) *tensor-field over E*.

Given our pair of smooth neural fields mapping from Euclidean space into the respective modality spaces, each mapping implicitly defines a Riemannian pull-back metric on the base space through the Jacobians ***J***_*ϕ*_ and ***J***_*ψ*_. The Riemannian pull-back metric is a classical approach in differential geometry for comparing manifolds in different spaces which may be expressed as maps from a common space.

While we offer a more formal description of this pull-back metric in Section A.1, this metric has a simple interpretation. Intuitively, *g*^*M*^ (*x*) (4) encodes the local anisotropy of modality/feature variation along different spatial directions in the modality *M*. A pair of points is deemed close if small spatial perturbations induce similar changes in the modality field. Thus, it “pulls” modality-specific structure back on the common Euclidean space. By mapping from Euclidean space into the modality spaces we are allowed to work in reference to the space common to two spatial multiomics datasets and compare distances in this common space.

For any pair of points *p, q* ∈ *M*, recall the *Riemannian (geodesic) distance* between them is given by the minimal length of a smooth curve *γ* : [0, 1] → *M* connecting them as measured in the metric

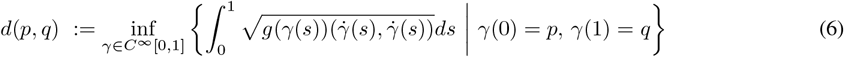

This coincides with the Euclidean distance when *M* is flat, and captures shortest paths in space curved according to the metric *g*. Now, given the pair of pull-back metrics for the two modalities defined on the common Euclidean space *E*

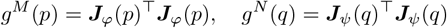

One may define Riemannian distances with respect to the pull-backs *g*^*M*^ and *g*^*N*^ for each modality (Definition 4). In particular, for each pair of spatial points *s, s*′ *E* in our first dataset and *t, t*′ ∈ *E* in our second, we compute geodesics *γ*^*M*^, *γ*^*N*^ in the modality-specific metrics *g*^*M*^, *g*^*N*^ and their Riemannian distance:

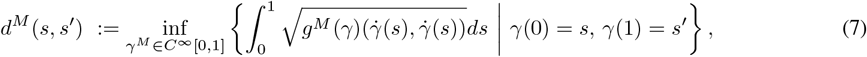

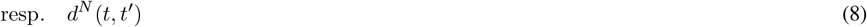

These paths *γ*^*M*^ and *γ*^*N*^ are highly intuitive (see, e.g. Figure 2), corresponding to a pull-back of the geodesics on each modality space onto *E* through the special requirement that *ϕ*(*E*) = ℳ and *ψ*(*E*) = 𝒩. Following training of the neural fields *ϕ, ψ*, computation of *g*^*M*^ and *g*^*N*^ at each point relies evaluation of the Jacobian, which is easily accomplished with automatic differentiation (3). However, (7) requires computing continuous, shortest-path geodesic curves. To approximate these, we simply discretize these paths at the resolution of the underlying spatial coordinate grid. In particular, given these metrics the geodesics are computed by building a *k*-NN graph 𝒢_*M*_ on the spatial points {*s*^*i*^}⊂ *E* with weights given by the local arc-length (infinitesimal distance unit) in the Riemannian metric between each neighboring pair of points. For edge (*i, i*′) one may define Δ_*ii*_ = (*s*^*i*^− *s*^*i*′^) and compute the local, symmetrized arc-length as

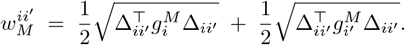

**Figure 2:**
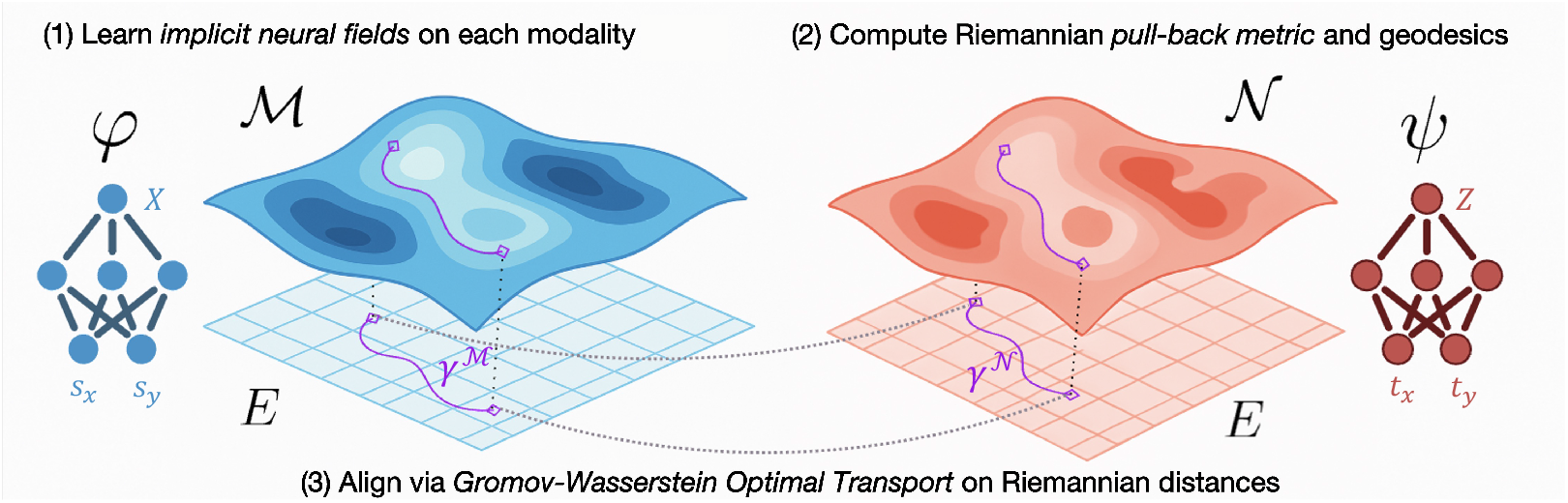
Manifold Gromov-Wasserstein (`MGW`). *ϕ, ψ* map from a Euclidean space *E* to distinct modality spaces ℳ and 𝒩. The maps *ϕ, ψ* define Riemannian pull-back metrics *g*^*M*^ and *g*^*N*^ through their Jacobians, from which one can compute geodesics *γ*^ℳ^, *γ*^𝒩^. Gromov-Wasserstein optimal transport yields an optimal alignment with respect to the Riemannian geodesic distances.

One repeats this for the second modality space 𝒩 and constructs 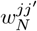 with graph 𝒢_*N*_ on the spatial coordinates {*t*^*j*^} of the second dataset. Now, with both spatial coordinates (vertices) and arc-lengths in the metric (edge weights), the Riemannian geodesic distances (7) are easily computed with standard routines for all-pairs shortest paths (APSP). Graph-based approximation of Riemannian geodesics on a *k*-NN graph is standard in manifold-learning and geometric graph methods (52; 8), and scales well to typical spatial omics resolutions.

### 2.2 Alignment with Manifold Gromov-Wasserstein

We begin by recalling the Wasserstein formulation of optimal transport (25). Let {*x*^1^, · · ·, *x*^*n*^ } and {*y*^1^, · · ·, *y*^*m*^ } be two datasets, and denote Δ_*k*_ to be the probability simplex of size *k*. In optimal transport, one encodes the two datasets as probability measures 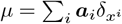 and 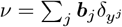 with probability vectors ***a*** ∈ Δ_*n*_, ***b*** ∈ Δ_*m*_ representing the weight of each point (e.g. uniform). Then, define the set of couplings with marginals ***a*** and ***b*** to be

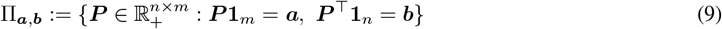

Given a cost matrix of distances between the points *X*^*i*^, *Y* ^*j*^, ***C***_*ij*_ := *c*(*X*^*i*^, *Y* ^*j*^), Wasserstein optimal transport (25) seeks a coupling ***P*** of minimal cost with respect to *c*

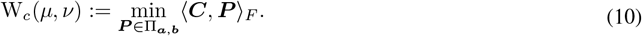

Where ⟨**A, B**⟩ _*F*_ = ∑_*ij*_ **A**_*ij*_**B**_*ij*_ denotes the Frobenius (matrix) inner product. While the Wasserstein formulation finds a coupling or alignment minimizing the distances between two points in a common space, often one seeks to compare {*X*^1^,, *X*^*n*^} ⊂ 𝒳 and {*Y* ^1^, · · ·, *Y* ^*m*^}⊂ 𝒴 in distinct metric spaces 𝒳, 𝒴. *Gromov-Wasserstein* optimal transport (32; 33) addresses this by allowing two cost functions *c*_1_ : 𝒳 × 𝒳 → ℝ_+_ and *c*_2_ : 𝒴 × 𝒴 → ℝ_+_, and instead comparing the metric distortion under ***P***

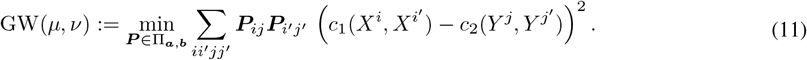

The Fused Gromov-Wasserstein (FGW) objective function (55) is a convex combination of the Wasserstein 10 and Gromov-Wasserstein 11 objectives with hyperparameter *α* ∈ (0, 1).

We define the *Manifold (Pull-Back) Gromov-Wasserstein Problem* to be the optimal coupling for (11) given the Riemannian pull-back distances 7 *d*^*M*^ (·, ·) and *d*^*N*^ (·, ·) of each modality onto the common base *E*.

#### Problem 2(Manifold Pull-Back GW Problem.).

*Given pull-back distances* (7) *d*^*M*^, *d*^*N*^ *and distributions* ***a*** ∈ Δ_*n*_ *over* 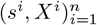 *and* ***b*** ∈ Δ_*m*_ *over* 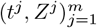, *Problem 2 is the optimization*

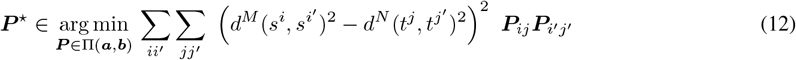

We highlight a few fundamental invariance and consistency properties of the `MGW` formulation.

**(P1)** (Consistency with Spatial GW) If the neural fields *ϕ* and *ψ* are identities; i.e. *ϕ*(*s*) = *s* and *ψ*(*t*) = *t*, then

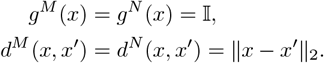

In the absence of information about the feature space, this offers an Occam’s Razor: (12) reduces exactly to the standard Gromov-Wasserstein problem on space *E*.

**(P2)** (Spatial Isometry Invariance, Proposition 1) Problem 2 is invariant to arbitrary translations *b* ∈ ℝ^*k*^, and orthogonal transformations *Q* ∈ 𝒪_*k*_ = {*Q* ∈ ℝ^*k*×*k*^ : *Q*^*T*^*Q* = *QQ*^*T*^ = **I**_*k*_}, so that solving 2 on *s*^*i*^ is equivalent to solving on

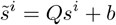

**(P3)** (Feature Isometry and Scaling Invariance, Proposition 2) Let *b, b*′ ∈ ℳ 𝒩, be any two constant vectors, let *λ* ℝ : *λ* ≠ 0 be a scaling, and *Q* ∈ 𝒪_*d*_, *U* ∈ 𝒪_*p*_ any two global orthogonal feature transformations. Then, the solution to Problem 2 is invariant to transformations of the feature space of the form

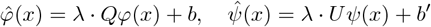

Notably, `MGW` is invariant to the coordinate representation or parameterization of both the feature and spatial spaces. Even when trained on differing neural architectures or parametrizations of the feature and spatial space, these guarantees hold and ensure `MGW` behaves robustly. This is in contrast to coordinate-dependent formulations such as (53; 7).

Manifold-GW offers a geometric unification for spatial multiomics alignment: for distinct feature modalities and a shared base space, all comparisons factor through a single term on this base – without the need for trade-off hyperparameters like fused Gromov-Wasserstein. By using a mapping *ϕ, ψ* to pull the modalities back to a shared space, the distances *d*^*M*^ (*s*^*i*^, *s*^*j*^) and *d*^*N*^ (*t*^*i*′^, *t*^*j*′^) are over a common space *E* and are compatible for comparison in a sense that raw feature distances, *c*_*M*_ (*X*^*i*^, *X*^*j*^) and *c*_*N*_ (*Z*^*i*^, *Z*^*j*^) may not be. Thus, even when ℳ ≠ 𝒩 are different spaces (e.g. representing distinct modalities), we are able to map directly between them by taking advantage of the product structure between the Euclidean space *E* and the pair ℳ, 𝒩.

We offer Algorithm 1 for `MGW` below, with an extended Algorithm S1 explicating all implementation details.

#### Algorithm 1

(Manifold GW: Geodesic Alignment).

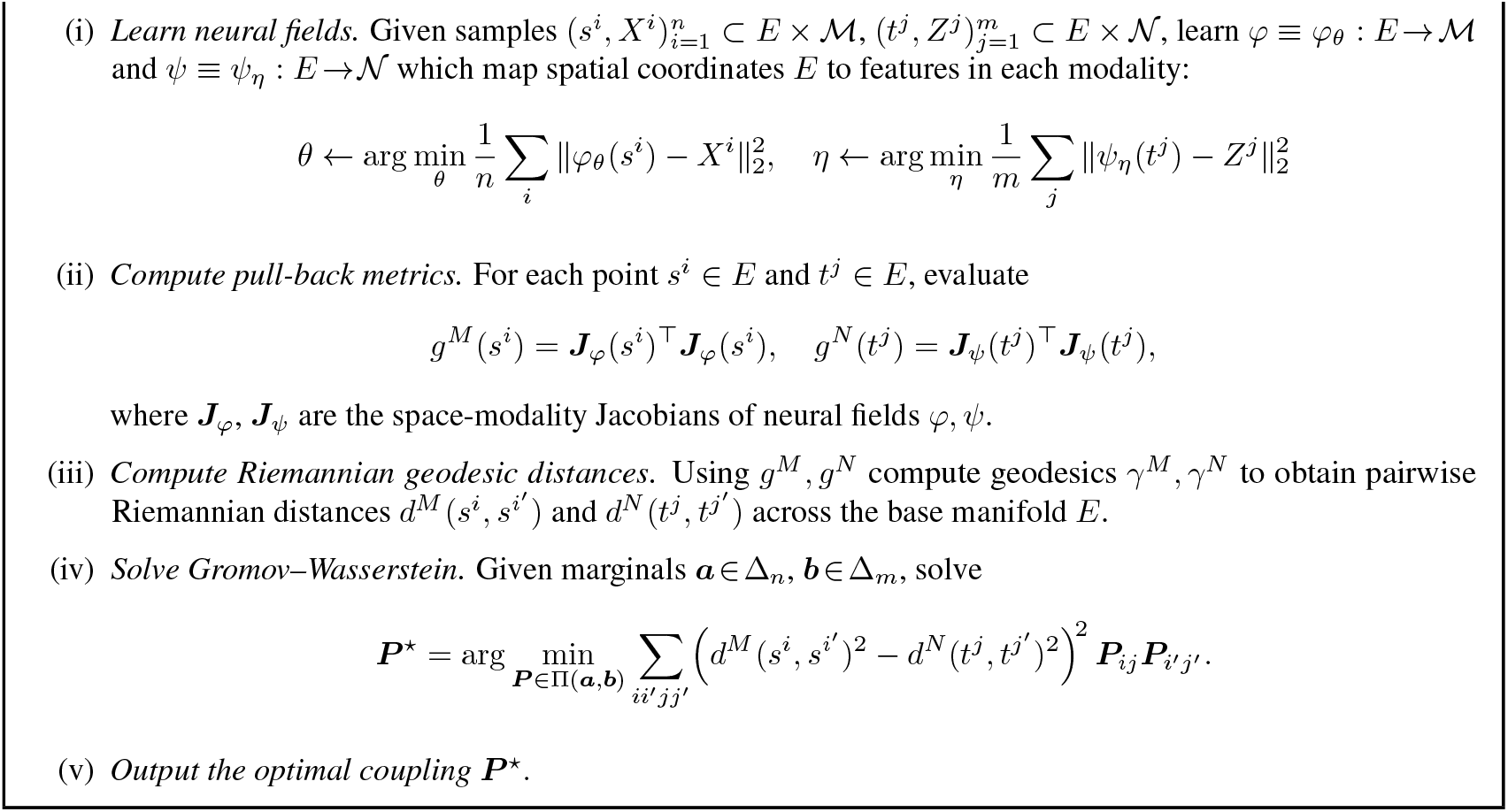

### 2.3 On Identifying the Alignment-Informative Feature Set

Observe that in 2 one assumes that the two spaces ℳ and 𝒩 exhibit joint structure. Yet the feature spaces of distinct modalities may not fully align or exhibit meaningful joint structure. In particular, there may be “marginal” sub-spaces within a modality which may be entirely independent of any features in the other modality, and there may be “joint” sub-spaces which exhibit high-correlation between the modalities. Thus, a natural pre-processing is to filter for such features 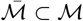 and 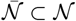 so that 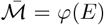 and 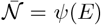 have high mutual information across *s* ∈ *E*. In Section C.6 we outline a pre-processing step to accomplish this by using CCA (*canonical correlation analysis* (20)) to filter for the subset of features across modalities with joint-structure prior to inferring the Riemannian metric with `MGW`.

## 3 Results

### 3.1 Spatiotemporal Transcriptomics of Mouse-Embryo

As an initial baseline and ablation of `MGW`, we benchmark on unimodal transcriptomics-transcriptomics data from consecutive time-points E9.5-10.5, E10.5-11.5, E11.5-12.5, and E12.5-13.5 of the mouse embryo using the Stereo-Seq platform (5). To mimic a multi-modal alignment and to isolate the impact of the proposed `MGW` cost, we focus the evaluation against standard Gromov-Wasserstein (GW) baselines: GW on Euclidean spatial distances, GW on Euclidean feature distances, unbalanced GW on feature distances with `moscot` TranslationProblem (26), and unbalanced GW on raw (Euclidean) feature geodesic distances with `SCOTv2` (10).

We assess alignments along two axes of quality: a metric of spatial realism, and a metric of feature alignment (Figure 3a). For the spatial accuracy of the alignment, we compute the migration metric (59; 30; 17): the average displacement of a cell when mapped into the coordinate frame of the target slice. For interpretability, we report the migration as percentage of the maximal extent of the slide. For the biological accuracy of the mapping, we compute the adjusted mutual information (AMI) of the predicted cell-types under the coupling against the ground-truth cell-type annotations present in (5).

**Figure 3:**
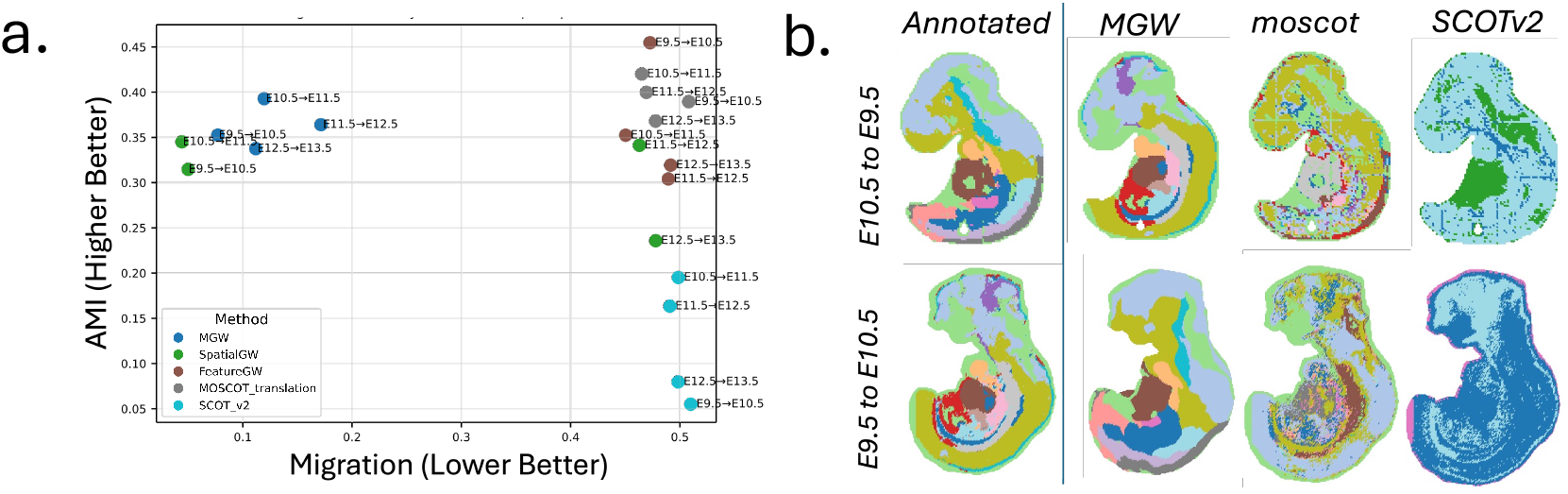
Comparison of Gromov-Wasserstein formulations on Stereo-Seq mouse-embryo timepairs E9.5-13.5 (5). (**a**) Comparison of alignments of pairs E9.5-10.5, E10.5-11.5, E11.5-12.5, and E12.5-13.5 in terms of migration distance and adjusted mutual information (AMI) with respect to published cell types. (**b**) Visualization of the most-likely cell-type (50% confidence threshold) predicted in the forward time direction (E9.5→E10.5) and backward time direction (E10.5→E9.5) transferred through each alignment.

We find that `MGW` achieves the best balance between spatial realism and feature alignment (Figure 3a). `MGW` has expected cell-migration of 7.7, 12.0, 17.2, and 11.2 % of the slide compared to GW on spatial distances alone: 5.0%, 4.4%, 46.3%, and 47.8%. Moreover, `MGW` achieves higher AMI values than spatial GW for all pairs: 0.353, 0.393, 0.364, 0.338 versus 0.315, 0.345, 0.341, 0.236. Notably, for the latter time pairs (E11.5-12.5 and E12.5-13.5) `MGW` achieves even lower migration distances than spatial-only GW – in these cases, the non-convex spatial GW optimization can converge to a global flip relative to the ground-truth due to nearly spatially symmetric coordinates (Figure S10). By learning the metric for each modality, `MGW` can break such symmetries. Relative to performing GW on features alone, which yields unrealistically large migrations of 47.3, 45.0, 48.9, and 49.2 % of the slide, `MGW` achieves much lower migration while maintaining comparable AMI values (0.353, 0.393, 0.364, 0.338 versus 0.455, 0.352, 0.304, 0.319). Likewise, compared to `moscot`, which produces migration distances of 50.8%, 46.5%, 46.9%, and 47.8% with AMI scores of 0.389, 0.420, 0.400, and 0.368, and to `SCOTv2`, which exhibits migrations of 51.0%, 49.9%, 49.1%, and 49.8% with markedly lower AMI (0.055, 0.195, 0.163, and 0.080), `MGW` attains minimal spatial displacement while maintaining high biological coherence, as shown in Figure 3b which illustrates the predicted cell-types at ≥ 50% confidence. These results highlight the importance of spatial information to correctly align spatially structured tissues and underscore the value of aligning the intrinsic manifold as opposed to raw expression or spatial coordinates.

### 3.2 Visium and Xenium Alignment of Human Colorectal Cancer

We perform an alignment of two multi-modal sections of a 10x Genomics colorectal cancer (CRC) dataset from the same donor (Sample P2 CRC) (37). This dataset includes a Visium CytAssist v2 section and a Xenium spatial transcriptomics section profiling the same tissue. While both datasets measure gene expression, they represent distinct measurement types: in Visium one performs sequencing of the full transcriptome (18k genes), while Xenium performs FISH imaging on a smaller panel of 422 genes. We benchmark 8 techniques on this dataset: `MGW`, Spatial GW, Feature GW, `moscot` (26), `SCOT` (12), `SCOTv2` (10), `PASTE2` (30), and POT (15). We measure alignment quality through mean expression cosine similarity between common Xenium-Visium genes, and (2) the spatial migration distance between aligned spots. Strong alignments maintain high cosine similarity in aligned cells, while also minimizing spatial distortion and crossings in the alignment.

We find that `MGW` achieves the best balance with high cosine similarity (mean 0.671 and median 0.689) and low migration distance (average 1.4% of the slide) (Figure 4, Table 4). Most of the other techniques – including Feature GW, `moscot`, `SCOT`, and `SCOTv2`– align based on feature-information alone and ignore the constraint of spatial context. As a result, they exhibit higher mean cosine scores 0.744/0.742/0.665/0.714, respectively, but also a highly unrealistic amount of spatial migration of 23.9 %/23.4%/23.4%/24.8%, respectively, of the spatial extent of the slide for the cell-cell mapping (Figure 4a). As `PASTE2` is not a multi-modal alignment method, it only uses spatial information and achieves a more realistic quantity of migration of only 0.1% of the slide but also a low expression cosine score of 0.437 indicating poor alignment of the transcription. Thus, `MGW` is the only technique with cosine-similarity at the level of feature-only methods and spatial distortion at the level of spatial-only methods.

**Figure 4:**
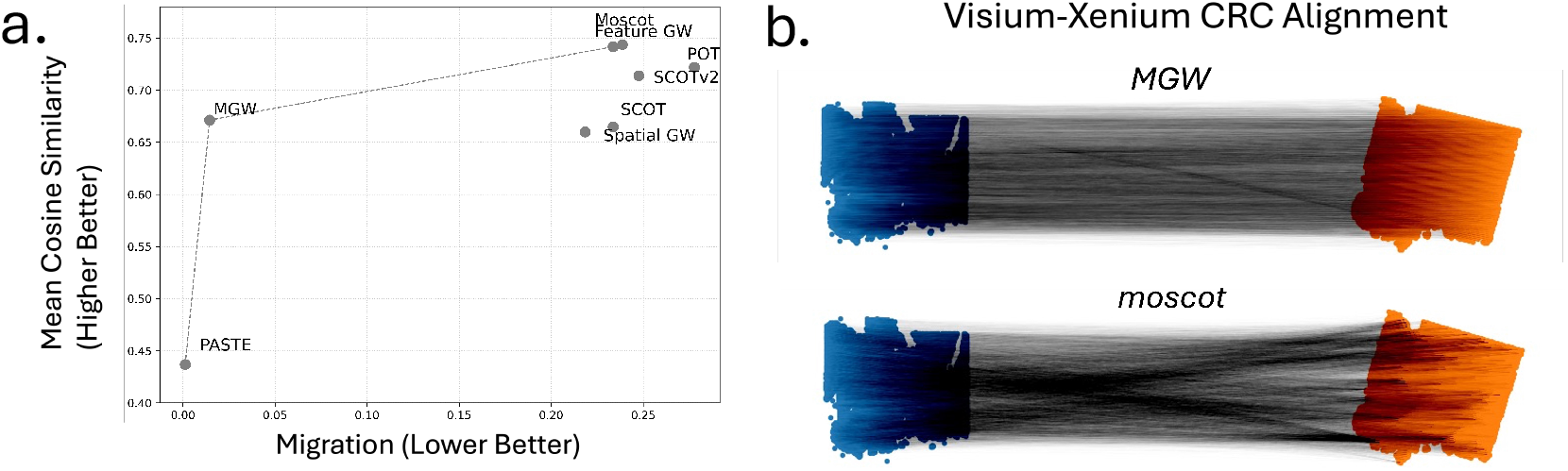
Comparison on Visium and Xenium datasets of colorectal cancer (37). (**a**) Cosine similarity and normalized migration of alignments on CRC Visium-Xenium alignment and (**b**) Visualization of `MGW` and `moscot TranslationProblem` alignment.

### 3.3 Metabolomics and Transcriptomics Alignment of Human Clear Cell Renal Cell Carcinoma (ccRCC)

We evaluated `MGW` on two tissue slices from clear-cell renal cell carcinoma (ccRCC), one assayed with a AFADESI-MSI spatial metabolomics and the other with 10x Genomics Visium spatial transcriptomics (dataset `Y7_T` from (22)). We benchmark on the task introduced in (53), which consists of: (1) aligning the multi-modal spatial metabolomic-transcriptomic datasets, (2) projecting the metabolomic profiles onto the transcriptomic coordinates through the alignment, and (3) computing joint-embeddings with the variational autoencoder (VAE) architecture of `SpatialMeta` (53). We compare `MGW` to multi-modal single-cell optimal-transport (OT) based techniques `SCOT` (11), `SCOTv2` (10), and `moscot` Translation (26), and to the non-OT based spatial metabolomics-transcriptomics method `SpatialMeta` (53), which introduced an Alignment Module based on `STAlign` (7) applied to the spatial coordinates. We also include Spatial-only GW and Feature-only GW as baselines. We ensure the training and architecture of the VAE are identical for all methods and repeat the entire evaluation pipeline, including alignment and training, across random seeds to ensure consistency. A complete summary of all metrics is provided in Table 2, and further details on our evaluation procedure and implementation may be consulted in Section C.

We find `MGW` reported the highest average spatial coherence across all methods, as quantified by Moran’s I across both modalities (0.6845 *±* 0.0134) (Figure 5c). Moran’s I assesses whether the top spatial transcriptomics and spatial metabolomics marker features selected from the Leiden clusters exhibit spatially smooth, region-specific expression patterns. `MGW` substantially outperforms `SpatialMeta` (0.5478), which is specifically designed for spatial transcriptomic and metabolomic alignment.

**Figure 5:**
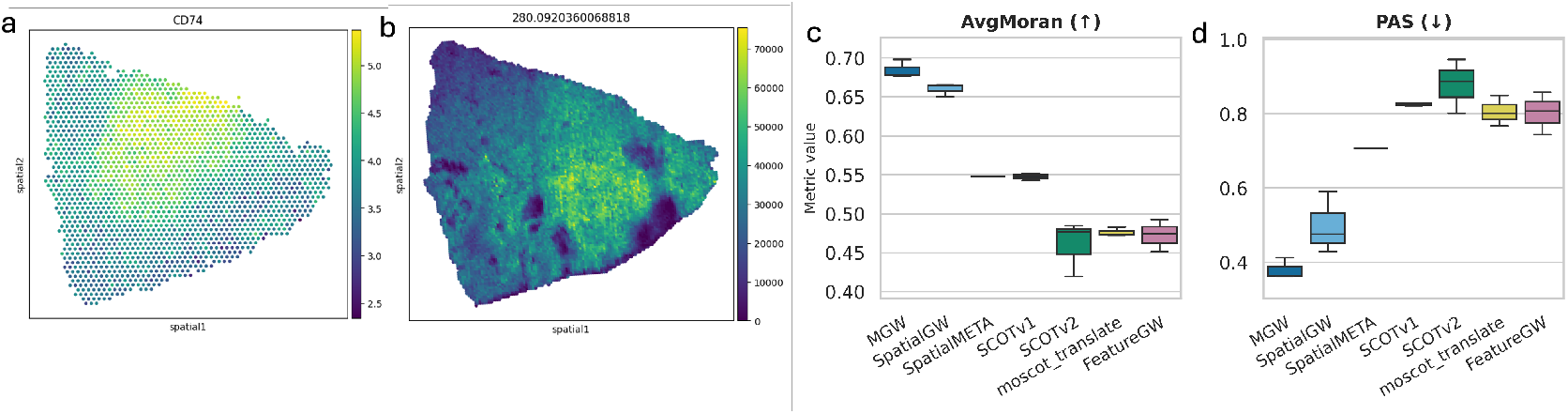
Cross-modal spatial transcriptomics and metabolomics alignment evaluation of `MGW`, spatial transcriptomics-metabolomics method `SpatialMeta`, and other OT baselines on the `Y7_T` slice from the ccRCC dataset (22). **(a)** The spatial distribution of gene *CD74* and **(b)** an associated metabolite with high Moran’s I. **(c)** Average Moran’s I on the two modalities and **(d)** `PAS` reported across three random seeds

`MGW` also outperforms the other methods on all three spatial continuity metrics used in (53) (Table 2). For example on `PAS` score, which quantifies how frequently a spot has neighbors belonging to a different cluster (see Section C), `MGW` achieved the lowest (best) value (Figure 5d), averaging 0.3797 across seeds, exceeding `SpatialMeta` (0.7063) as well as `SCOT`, `SCOTv2` and `moscot` with scores of 0.8255, 0.8784, and 0.8058. In addition, the scores exceed those of Feature and Spatial GW (0.8029 and 0.4985, respectively). Techniques which aligned using spatial coordinates alone, such as `SpatialMeta` (which uses `STAlign`) and Spatial Gromov-Wasserstein, exhibited stronger performance on this task than ones relying on feature-information alone. This reflects the similarity of the aligned spatial grids (Figure 5a,b).

### 3.4 Spatial Metabolomics and Transcriptomics Alignment of Striatum in Human Brain

Lastly, we align two post-mortem sections from the striatum region of the human brain. One section contains spatial profiling of low molecular weight metabolites determined using (MALDI)-MSI mass-spectrometry, and the other section contains spatial profiling of mRNA transcripts measured using the 10x Genomics Visium technology (56). These sections are from a patient with Parkinson’s disease – a disorder characterized by dopamine depletion in the striatum (56). To evaluate `MGW` in comparison to other techniques, we assess the correspondence between the concentration of dopamine and its immediate breakdown products at various mass:charge ratios *m/z* in the spatial metabolomics dataset compared to the published annotations of Visium spots by dopamine-positive identity provided in (56) (Figure 6a). Specifically, (56) identifies 4 key dopamine metabolites, including DA Single (singly-derivatized dopamine) at *m/z* = 421.19, DA double (doubly-derivatized dopamine) at *m/z* = 674.28, the dopamine-breakdown product 3MT (3-Methoxytyramine) at *m/z* = 435.21, and the dopamine-breakdown product DOPAC double at *m/z* = 698.24. Dopamine-associated neurons were defined using the dopamine-high (Cd) annotations of the spatial transcriptomics slide (56).

**Figure 6:**
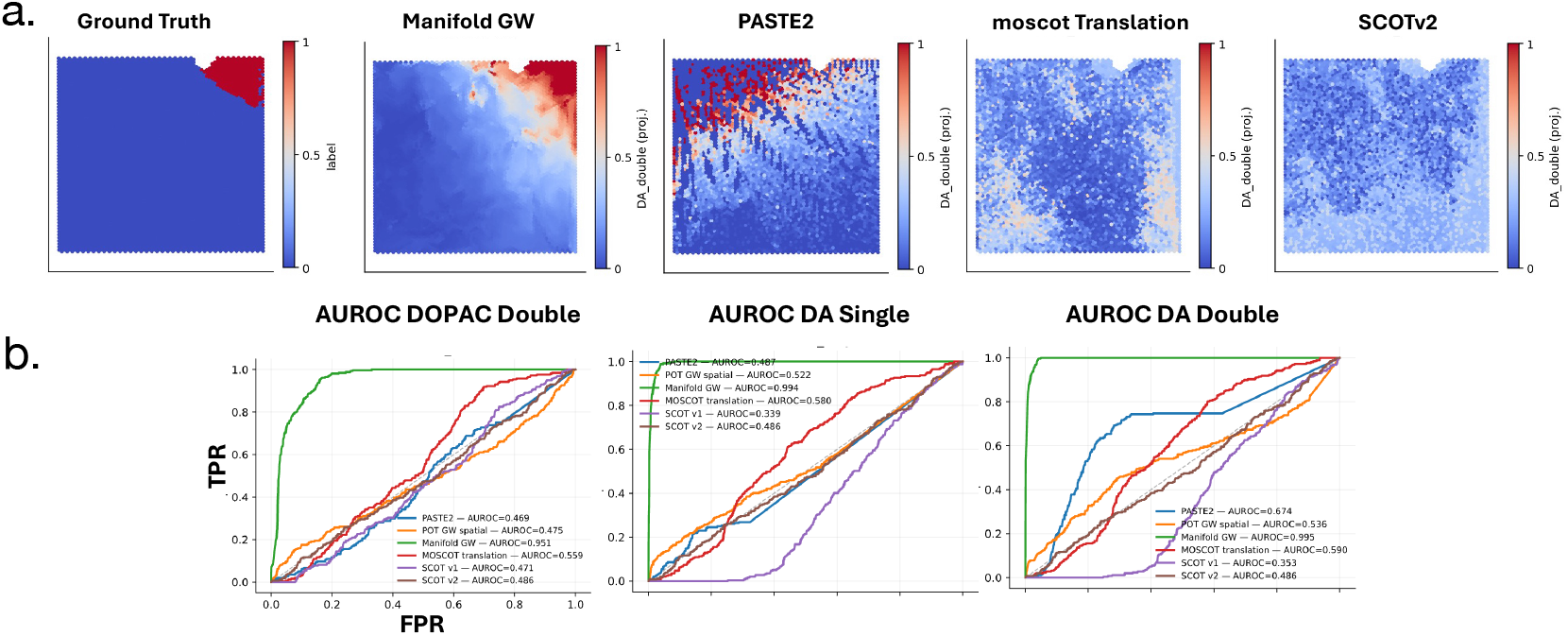
(**a**) (Left) Published (“Ground-truth”) dopamine labels on Visium slide (56). (Right) Normalized barycentric projection of doubly-derivatized dopamine onto Visium slide across methods. (**b**) AUROC curves for the overlap between the locations of Dopamine metabolites DA Single (singly-derivatized *m/z* = 421.19), DA double (doubly-derivatized *m/z* = 674.28), 3MT (dopamine-breakdown product 3-Methoxytyramine, *m/z* = 435.21), and DOPAC double (dopamine-breakdown product *m/z* = 698.24) and the annotated locations of dopamine-associated neurons on Visium spatial transcriptomics slide for each method.

We compared `MGW` to multi-modal single-cell alignment techniques `moscot` Translation (26), `SCOT` (11), and `SCOTv2` (10), as well as two spatial-only baselines: `PASTE2` with spatial information alone (30), and spatial-only GW. We choose the latter baselines in order to assess that the efficacy of the `MGW` alignment is not merely due to spatial information. Each method produces a coupling ***P*** between the locations on the metabolomics and spatial transcriptomics slices. We compute a barycentric projection (40) of the metabolite intensities through each ***P*** onto the ST coordinates. All methods except `PASTE2`, which performs partial alignment, transfer the full dopamine mass-intensity to the ST slide. We quantify alignment accuracy as as the overlap between projected dopamine intensities from the metabolomics slices and the annotated binary dopamine-high spot labels on the transcriptomics slice, and computed the AUROC and and AUPRC as a function of the predicted metabolite scores (Figure 6b). As the dopamine metabolites and dopamine-high spots form a weak, sharply localized signal (See Figure S8 for unscaled, raw-intensities), this benchmark assesses high-precision mapping of this small but biologically significant region. Low scores indicate failure to accurately map this local signal rather than poor global alignment.

`MGW` achieves the highest scores across all four metabolites (Table 1, Figure 6b). Specifically, the AUROC and AUPRC scores for 3MT, DA double, DA single, and DOPAC double were 0.991/0.846, 0.995/0.907, 0.994/0.882, and 0.951/0.455, respectively. These consistently exceeded the scores of other methods by large margins – for example, `moscot` Translation achieved 0.615/0.063, 0.590/0.058, 0.580/0.057, and 0.559/0.054, while the strongest spatial baseline `PASTE2` reached 0.667/0.084, 0.674/0.090, 0.487/0.055, and 0.469/0.046. The alignment of Spatial GW performed near chance (AUROC ~ 0.52). Visualizations of the projections (Figure 6a) corroborate the observed quantitative trend: `MGW` correctly projects the dopamine metabolites onto the ground-truth dopamine-annotated regions with high spatial localization and accuracy. Other methods yield diffuse or incorrectly localized patterns, indicating that accurate spatial multiomic alignment requires both spatial and feature modalities. In this high-precision, localized prediction task, `MGW` is the only method which projects dopamine metabolites to the dopamine-annotated neurons.

**Table 1:** Comparison of alignment methods across dopamine-related MSI targets. Best scores per target are shown in bold.

| Method | Target | $m/z$ | AUROC | AUPRC |
| --- | --- | --- | --- | --- |
| Manifold GW | 3MT | 435.21 | **0.991** | **0.846** |
| PASTE2 | 3MT | 435.21 | 0.667 | 0.084 |
| MOSCOT (translation) | 3MT | 435.21 | 0.615 | 0.063 |
| POT-GW (spatial) | 3MT | 435.21 | 0.522 | 0.081 |
| SCOT_v2 | 3MT | 435.21 | 0.486 | 0.051 |
| SCOT_v1 | 3MT | 435.21 | 0.380 | 0.038 |
| Manifold GW | DA_double | 674.28 | **0.995** | **0.907** |
| PASTE2 | DA_double | 674.28 | 0.674 | 0.090 |
| MOSCOT (translation) | DA_double | 674.28 | 0.590 | 0.058 |
| POT-GW (spatial) | DA_double | 674.28 | 0.536 | 0.085 |
| SCOT_v2 | DA_double | 674.28 | 0.486 | 0.051 |
| SCOT_v1 | DA_double | 674.28 | 0.353 | 0.037 |
| Manifold GW | DA_single | 421.19 | **0.994** | **0.882** |
| MOSCOT (translation) | DA_single | 421.19 | 0.580 | 0.057 |
| POT-GW (spatial) | DA_single | 421.19 | 0.522 | 0.081 |
| PASTE2 | DA_single | 421.19 | 0.487 | 0.055 |
| SCOT_v2 | DA_single | 421.19 | 0.486 | 0.051 |
| SCOT_v1 | DA_single | 421.19 | 0.339 | 0.036 |
| Manifold GW | DOPAC_double | 698.24 | **0.951** | **0.455** |
| MOSCOT (translation) | DOPAC_double | 698.24 | 0.559 | 0.054 |
| SCOT_v2 | DOPAC_double | 698.24 | 0.486 | 0.051 |
| POT-GW (spatial) | DOPAC_double | 698.24 | 0.475 | 0.061 |
| SCOT_v1 | DOPAC_double | 698.24 | 0.471 | 0.045 |
| PASTE2 | DOPAC_double | 698.24 | 0.469 | 0.046 |

**Table 2:**
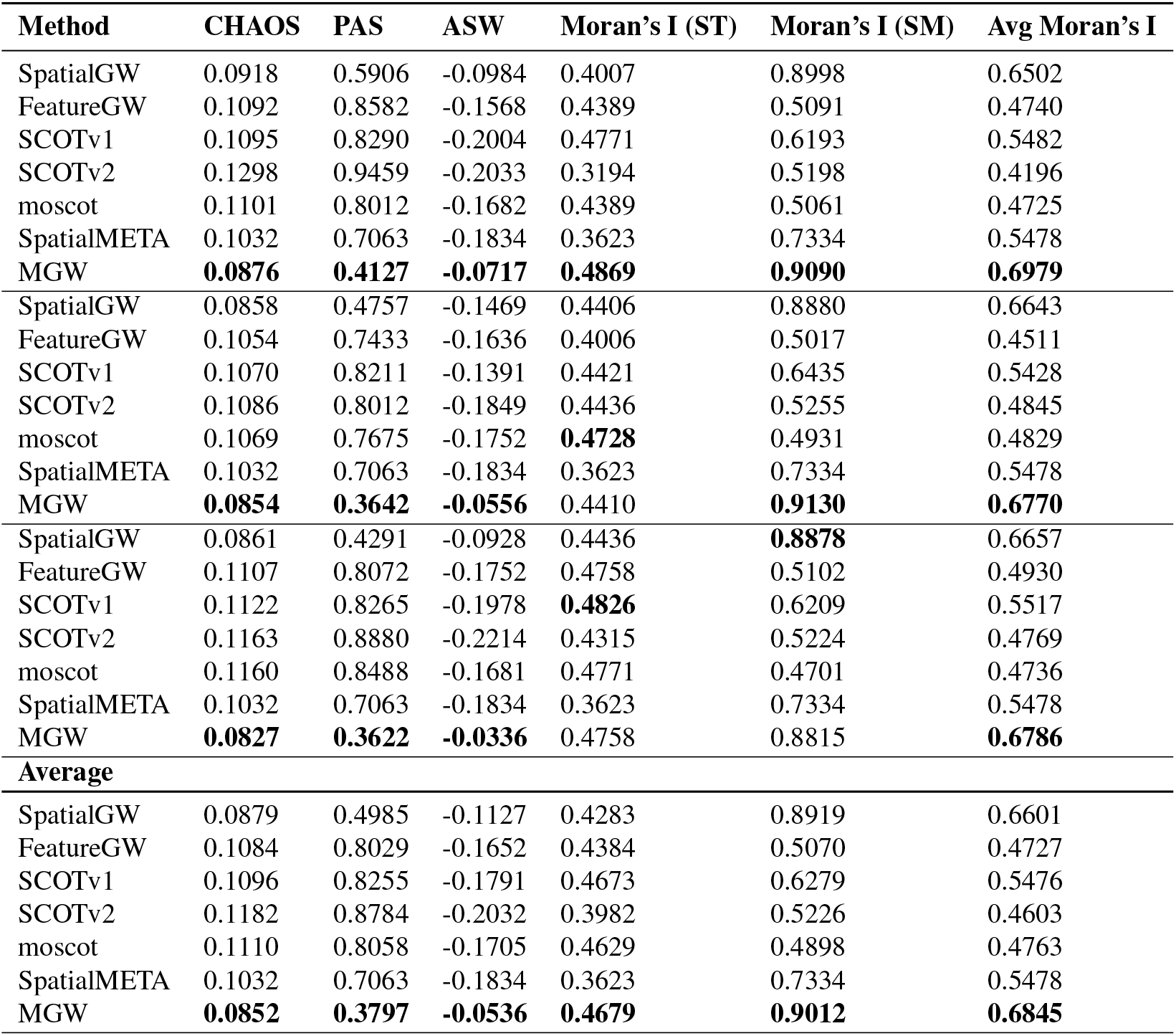
Comparison of alignment methods across metrics across three random seeds. Best scores per metric are shown in bold.

**Table 3:** Pairwise alignment performance across developmental stages. Lower migration and higher projected AMI indicate better cross-slice alignment quality.

| Pair | Method | Migration ↓ | Proj. AMI ($A \leftrightarrow B$) ↑ |
| --- | --- | --- | --- |
| E9.5→E10.5 | FeatureGW | 0.473 | 0.455 |
| E9.5→E10.5 | MGW | 0.077 | 0.353 |
| E9.5→E10.5 | MOSCOT_translation | 0.508 | 0.389 |
| E9.5→E10.5 | SCOT_v2 | 0.510 | 0.055 |
| E9.5→E10.5 | SpatialGW | 0.050 | 0.315 |
| E10.5→E11.5 | FeatureGW | 0.450 | 0.352 |
| E10.5→E11.5 | MGW | 0.120 | 0.393 |
| E10.5→E11.5 | MOSCOT_translation | 0.465 | 0.420 |
| E10.5→E11.5 | SCOT_v2 | 0.499 | 0.195 |
| E10.5→E11.5 | SpatialGW | 0.044 | 0.345 |
| E11.5→E12.5 | FeatureGW | 0.489 | 0.304 |
| E11.5→E12.5 | MGW | 0.172 | 0.364 |
| E11.5→E12.5 | MOSCOT_translation | 0.469 | 0.400 |
| E11.5→E12.5 | SCOT_v2 | 0.491 | 0.163 |
| E11.5→E12.5 | SpatialGW | 0.463 | 0.341 |
| E12.5→E13.5 | FeatureGW | 0.492 | 0.319 |
| E12.5→E13.5 | MGW | 0.112 | 0.338 |
| E12.5→E13.5 | MOSCOT_translation | 0.478 | 0.368 |
| E12.5→E13.5 | SCOT_v2 | 0.498 | 0.080 |
| E12.5→E13.5 | SpatialGW | 0.478 | 0.236 |

**Table 4:** Comparison of alignment methods on cosine similarity and migration metrics for Visium-Xenium alignment (CRC Colorectal Cancer tumor sample).

| Method | Mean Cosine | Median Cosine | Migration |
| --- | --- | --- | --- |
| MGW | 0.671 | 0.689 | 0.014 |
| Spatial GW | 0.660 | 0.681 | 0.218 |
| Feature GW | 0.744 | 0.753 | 0.239 |
| Moscot | 0.742 | 0.754 | 0.234 |
| SCOT | 0.665 | 0.694 | 0.234 |
| SCOTv2 | 0.714 | 0.725 | 0.248 |
| PASTE | 0.437 | 0.455 | 0.001 |
| POT | 0.722 | 0.738 | 0.278 |

## 4 Discussion

We introduce Manifold Gromov-Wasserstein (`MGW`) to align spatial multiomic data across heterogenous feature spaces. `MGW` assumes these spaces arise as the image of smooth maps from a shared Euclidean domain and learns modality-specific Riemannian metrics via neural fields. By using these Riemannian pull-back metrics, `MGW` compares the intrinsic structure of these modalities on their common spatial base, avoiding any hyperparameters balancing spatial and feature similarity. We demonstrate Manifold Gromov-Wasserstein on a diverse set of modalities – Visium and Stereo-seq sequencing, Xenium imaging, and MALDI-MSI and AFADESI-MSI mass-spectrometry metabolomics – highlighting the generality of `MGW`.

Several limitations suggest directions for future work. First, as `MGW` relies on two neural networks *ϕ, ψ* to learn the underlying cost for each modality space, the choice of underlying architecture may improve the quality of metric-learning and the alignment itself. While we focus on lightweight MLPs, a natural question is how the class of neural network may impact performance. Second, spatial multiomic data often contain modality-specific features with no shared structure. While we propose a pre-processing step based on CCA to mitigate this (Section C.6), we believe identifying mutually informative feature subspaces would improve estimation of the metric. Third, optimal transport variants such as unbalanced (46; 26; 10), semi-relaxed (17), and partial (30) OT often improve alignment robustness. Although `MGW` supports unbalanced marginals, we used balanced OT in our experiments to highlight the effect of the learned Manifold-GW cost.

Beyond spatial multiomic alignment, Riemannian distances may be especially advantageous for highly non-linear transformations such as spatiotemporal transcriptomics, where intrinsic manifold geometry is more informative than raw Euclidean distances (Section 3.1). Our ablations additionally indicate that the `MGW` Riemannian distances constitute a stronger GW term than either spatial or feature distances alone, suggesting FGW style frameworks may also benefit from adopting this cost.

Manifold Gromov-Wasserstein offers a principled way to align heterogeneous spatial omics datasets through deep Riemannian metric learning and optimal transport. More broadly, our results underscore that learning intrinsic geometry provides a powerful unifying principle for aligning nonlinear manifold data across the spatial multiomic universe.

## Acknowledgements

This research was supported by NIH/NCI grant U24CA248453 to B.J.R and by Ludwig Cancer Research.

## A Appendix

### A.1 Pull-back Metrics and Base-Space Parametrization

Suppose we are given two Riemannian manifolds (ℳ, *g*) and (𝒩, *h*) and measures (datasets) *µ* ∈ 𝒫(ℳ), *ν* ∈ 𝒫(𝒩) supported on each space: for instance, for single-cell transcriptomics ℳ could represent an embedded sub-manifold of the *d*-dimensional measurements of transcript features ℳ ⊂ ℝ^*d*^, and for a distinct modality, e.g. metabolomics, 𝒩 could represent a sub-manifold of the *p*-dimensional measurements of metabolites 𝒩 ⊂ ℝ^*p*^ where we may have that *p* ≠ *d* and dim(ℳ) ≠ dim(𝒩). In general, it may appear to be a formidable task to align such datasets across such incomparable spaces, with no “natural” notion of distance immediately evident.

However, suppose that we instead take measurements on the product space ℳ × *E* and 𝒩 × *E* for some common base manifold (*E, g*^*E*^). This may correspond to Euclidean space *E* = ℝ^2^ or *E* = ℝ^3^, e.g. when one reads the product of feature vectors with spatial coordinates as in spatial transcriptomics or spatial metabolomics. Recent, a remarkable line of work (6) have shown that one may learn an implicit neural representation of such product datasets. In particular, (6) shows that one may represent data (*E*, ℳ) as a smooth and differentiable map *φ* ∈ *C*^*k*^ from a base-space *E* into a feature space ℳ, *φ* : *E* → ℳ. In other words, given spatial coordinates *s*^*i*^ and product vector *p*^*i*^ = (*s*^*i*^, *X*^*i*^) ∈ *E* × ℳ one may instead represent *X*^*i*^ := *φ*(*s*^*i*^) and thus *p*^*i*^ = (id, *φ*) ◦ *s*^*i*^ for smooth, differentiable *φ*: a spatial transcriptomics dataset may be implicitly represented as a smooth function of its spatial coordinates. Thus, we state a first key assumption.

#### Assumption 1.

*Given* (ℳ, *g*) *and* (𝒩, *h*) *Riemannian, a common base manifold* (*E, g*^*E*^), *and points p*^*i*^ = (*s*^*i*^, *X*^*i*^) ∈ *E* × ℳ *and q*^*j*^ = (*s*^*j*^, *Z*^*j*^) ∈ *E* × 𝒩 *we assume there there exist two C*^*k*^ *local-parametrizations*

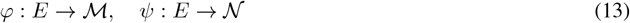

*which satisfy*

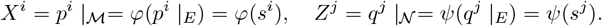

As in (6), we optimize for such a pair of differentiable mappings with a neural network encoding each dataset implicitly. One solves this pair of maps *φ, ψ* as

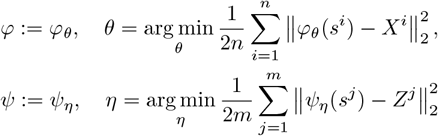

Given explicit maps *φ, ψ* from the base *E* into ℳ and 𝒩 with associated metrics on each space, there are natural vector fields defined through the mappings and a natural metric. In particular, for a vector field on *E, V* ∈ 𝔛 (*E*), one has associated push-forward vector fields on ℳ, 𝒩 given by

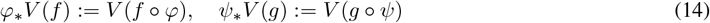

defined by 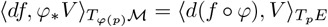 for any test function *f* ∈ *C*^*∞*^(ℳ) (resp. 𝒩). With a correspondence between spaces, we may measure manifold distances on each space with a *pull-back metric*.

#### Definition 3 (Pull-Back Metric.).

*Given manifolds* (*E, g*^*E*^), (𝒩, *h*) *and mapping ψ* : *E* → 𝒩, *the pull-back metric of h is the symmetric* (2, 0)*-tensor on E defined by*

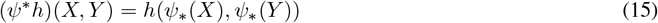

From the duality of 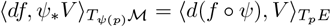, one identifies

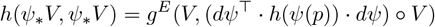

so that one may write this in terms of the Jacobian as

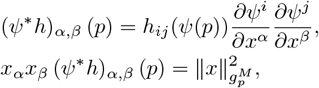

For metric 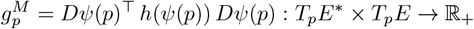 defined at any point *p* ∈ *E*.

Now, suppose for simplicity that *g* and *h* are taken to be the standard Euclidean metrics *g*_*αβ*_ = *δ*_*αβ*_ in their respective spaces (i.e. in normal coordinates). Then, given a non-trivial pair of mappings *φ, ψ* one can find the following pull-back of both ℳ, 𝒩 to the common Euclidean space *E*:

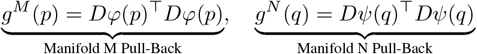

Thus, we arrive at a remarkable identification: if we measure in a product space *E* × ℳ and *E* × 𝒩, and the pull-back maps (*φ, ψ*) exist, one may pull manifold-distances back to a *common* base space. Observe that *g*^*M*^ and *g*^*N*^ are *tensor fields* over *E*: each coordinate *p, q* has its own metric tensor *g*^*M*^ (*p*) and *g*^*N*^ (*q*), which may vary signficantly over *E*.

### A.2 Metric Distances and Energy

Let us recall the fundamental notion of distance used with respect to a metric *g*. The infinitesimal unit *ds* is known as the line element and gives an infinitesimal unit of arc length (distance). To extend this to distances over curves, define any curve over the manifold by *γ* : [0, 1] → *E*, and the line-element of this curve at any time *t* and point *γ*(*t*) by

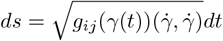

one measures the distance of a curve *γ* over time *t* with the arc-length integral

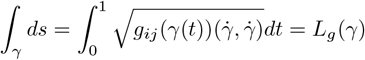

For local coordinates *X* = (*x*^1^, · · ·, *x*^*n*^) one defines the infinitesimal squared-distance in *g* by

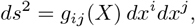

So that the extension for a curve *γ* is *ds* 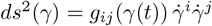. The *energy* of a curve *γ* is defined as the integral of this squared-distance unit along the curve *γ*

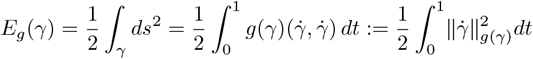

Suppose one normalizes *γ* to be parametrized by arc-length, i.e. the unique constant speed parametrization of *γ* so that 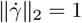_, then one may define a geodesic distance between any points *x*^*a*^, *x*^*b*^ by the minimal distance between these_ points over all curves *γ* connecting them:

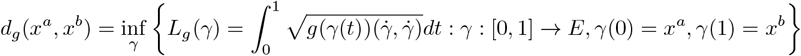

Which, for constant speed curves, is related to the energy by

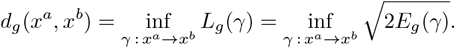

The argmin of these problems is the unique constant-speed *γ*^*∗*^ from *x*^*a*^ to *x*^*b*^, and the minimum value of *d*_*g*_(*x*^*a*^, *x*^*b*^) is referred to as either a Riemannian distance or a geodesic distance between the pair of points. Thus, given a potentially point-dependent metric *g* on ℳ, one may define a geodesic distance between any pair of points. In the standard Euclidean case, the definitions above automatically recover the standard Euclidean distance and distance-squared, where *g*_*ab*_(*x*) = *g*_*ab*_ := *δ*_*ab*_ reduces to a trivial constant metric which lacks any dependence on the location of *x* on ℳ.

### A.3 Manifold Alignment via Geodesic Pull-Back Distances

Using the notions of distance above, we provide a formulation for the Gromov-Wasserstein problem (Definition 5) which offers a geometric unification of the alignment problem between spaces using the pull-back. In particular, this cost relies on geodesic pull-back distances under the pull-back metrics *g*^*M*^, *g*^*N*^ over the common base *E*.

#### Definition 4 (Riemannian Pull-Back Distances.).

*Suppose we are given two feature manifolds* (ℳ, *g*), (𝒩, *h*), *a common base manifold* (*E, g*^*E*^), *and points p*^*i*^ = (*s*^*i*^, *X*^*i*^) ∈ *E* × ℳ *and q*^*j*^ = (*s*^*j*^, *Z*^*j*^) ∈ *E* × 𝒩. *Further, suppose that Assumption 1 holds with associated maps φ* : *E* → ℳ, *ψ* : *E* → 𝒩. *We define the M-distance and N-distance of any two points x*^*s*^, *x*^*t*^ ∈ *E to be the associated geodesic pull-back distances:*

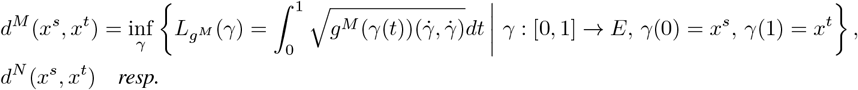

*For pull-back metrics g*^*M*^, *g*^*N*^ *defined by*

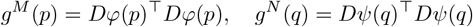

Now, given appropriate notions of pairwise distance in the common Euclidean space *E*, we may define the associated Gromov-Wasserstein (GW) problem (32; 33) as the natural notion of metric distortion between two spaces with two metrics.

#### Definition 5 (Manifold (Pull-back) Gromov-Wasserstein.).

*Given Riemannian pull-back distances d*^*M*^ (·, ·) *and d*^*N*^ (·, ·) *(Definition 4), define the associated squared distortions to be C*^*M*^ (·, ·) := *d*^*M*^ (·, ·)^2^, *and C*^*N*^ (·, ·) := *d*^*N*^ (·, ·)^2^. *For two distributions µ*^*M*^ ∈ 𝒫(*E*) *and µ*^*N*^ ∈ 𝒫(*E*), *the manifold pull-back GW Distance between µ*^*M*^ *and µ*^*N*^, *is*

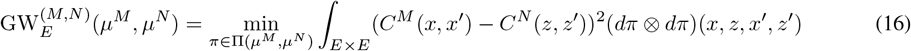

In other words, this is the standard Gromov-Wasserstein distance with Riemannian distance matrices built from Definition 4. Written explicitly in full, this is equal to

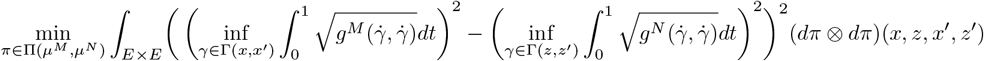

The corresponding discrete problem for Definition 4 is given for two datasets of points 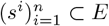 and 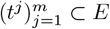 in the Euclidean base-space by

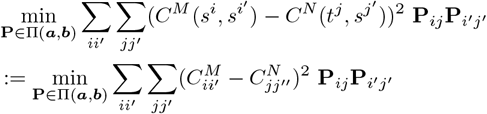

We summarize this discrete formulation in Problem 2. In the next section, we discuss a number of desirable properties and invariances of Definition 5 which makes it a natural option for aligning data that admits a product structure across two spaces.

### B Properties of the Gromov-Wasserstein Distance on the Metric Pull-back

#### B.1 Consistency under identity mapping

Observe that if *φ, ψ* := id so that *φ*(*s*), *ψ*(*t*) := *s, t* – i.e., there is no feature modality, one has that

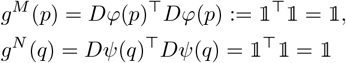

so that the metric tensor is the identity, and thus the geodesic distances become Euclidean distances, *d*^*M*^ (*x*^*s*^, *x*^*t*^) = ‖ *x*^*s*^ − *x*^*t*^ ‖_2_. As a consequence, (16) simply reduces to a Gromov-Wasserstein problem on the raw spatial distances. Thus, (16) is consistent: in the case that no feature modality is present it recovers standard Gromov-Wasserstein on the spatial distances alone.

#### B.2 Spatial Invariances

##### Proposition 1 (Spatial Invariances of Manifold Gromov-Wasserstein.).

*Problem 2 is invariant to arbitrary translations b* ∈ ℝ^*k*^, *and orthogonal transformations Q* ∈ 𝒪_*k*_ = {*Q* ∈ ℝ^*k*×*k*^ : *Q*^⊤^ *Q* = *QQ*^⊤^ = **I**_*k*_}, *so that solving 2 on s*^*i*^ *is equivalent to solving on*

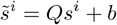

*Proof*. Let *g*^*N*^ = *Dφ*^⊤^*Dφ* on a Euclidean manifold *E* = ℝ^*k*^. To assess spatial invariances, we consider which maps Φ : *E* → *E* applied to the points *x* → Φ(*x*) maintain the solution of the objective. Let us consider the action of the class of rigid-body transformations for arbitrary translations *b* ∈ ℝ^*k*^, and orthogonal transformation *Q* ∈ ℝ^*k*×*k*^ : *Q*^⊤^*Q* = *QQ*^⊤^ = **I**_*k*_. Suppose one learns the maps 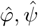 over a rigid-body transformation of the coordinates *y* = *Qx* + *b, x* = *Q*^⊤^(*y* − *b*) and denote the maps over the original coordinates by *φ, ψ*. Since the features are identical under rotation of the spatial grid, one has that within the compact set Ω ⊂ *E* our points are supported on:

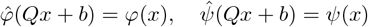

and thus for the modified metric tensor *ĝ*^*M*^ one identifies

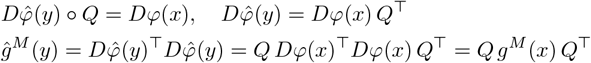

implying that for any curve *γ* and its rigid transformation *γ*_*∗*_ = *Qγ* + *b* one has 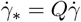 so that

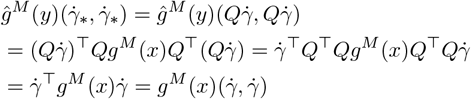

So that infinitesimal line elements coincide, and

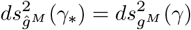

and integrating along curves yields that the geodesic distance in *M* (resp. *N*) is invariant under rigid motions

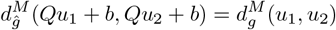

#### B.3 Feature Invariances

Similarly to the spatial case, for the feature case we evaluate which transformations *T* ^*φ*^, *T* ^*ψ*^ of the feature maps *φ, ψ* maintain the modified objective.

##### Proposition 2 (Feature Invariances of Manifold Gromov-Wasserstein.).

*Let b, b*^*′*^ ∈ ℳ, 𝒩 *be any two constant vectors, let λ* ∈ ℝ : *λ* ≠ 0 *be a scaling, and Q* ∈ 𝒪_*d*_, *U* ∈ 𝒪_*p*_ *any two global orthogonal feature transformations. Then, the solution to Problem 2 is invariant to transformations of the feature space of the form*

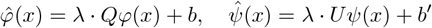

*Proof*. Let us focus on the following class of transformations for any constant vector *b, b*^*′*^ ∈ ℳ, 𝒩, a scaling *λ* ∈ ℝ : *λ* ≠ 0, and a global orthogonal feature transformation *Q* ∈ 𝒪_*d*_, *U* ∈ 𝒪_*p*_

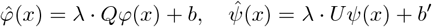

One may verify that the Jacobian is necessarily translation-invariant, yielding

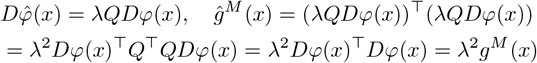

With infinitesimal line-elements differing by a homogeneous constant, and implying

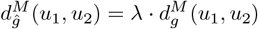

So that, applying the same reasoning to 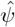, we have

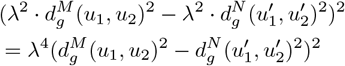

And have 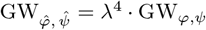 that the solution is unchanged to this class of transformation and is conformally invariant to the change of metric *g* → *λ*^2^*g*.

##### Algorithm 1

Manifold GW (M-GW): Geodesic Distances via a pull-back metric on the common base *E* (Long Version)

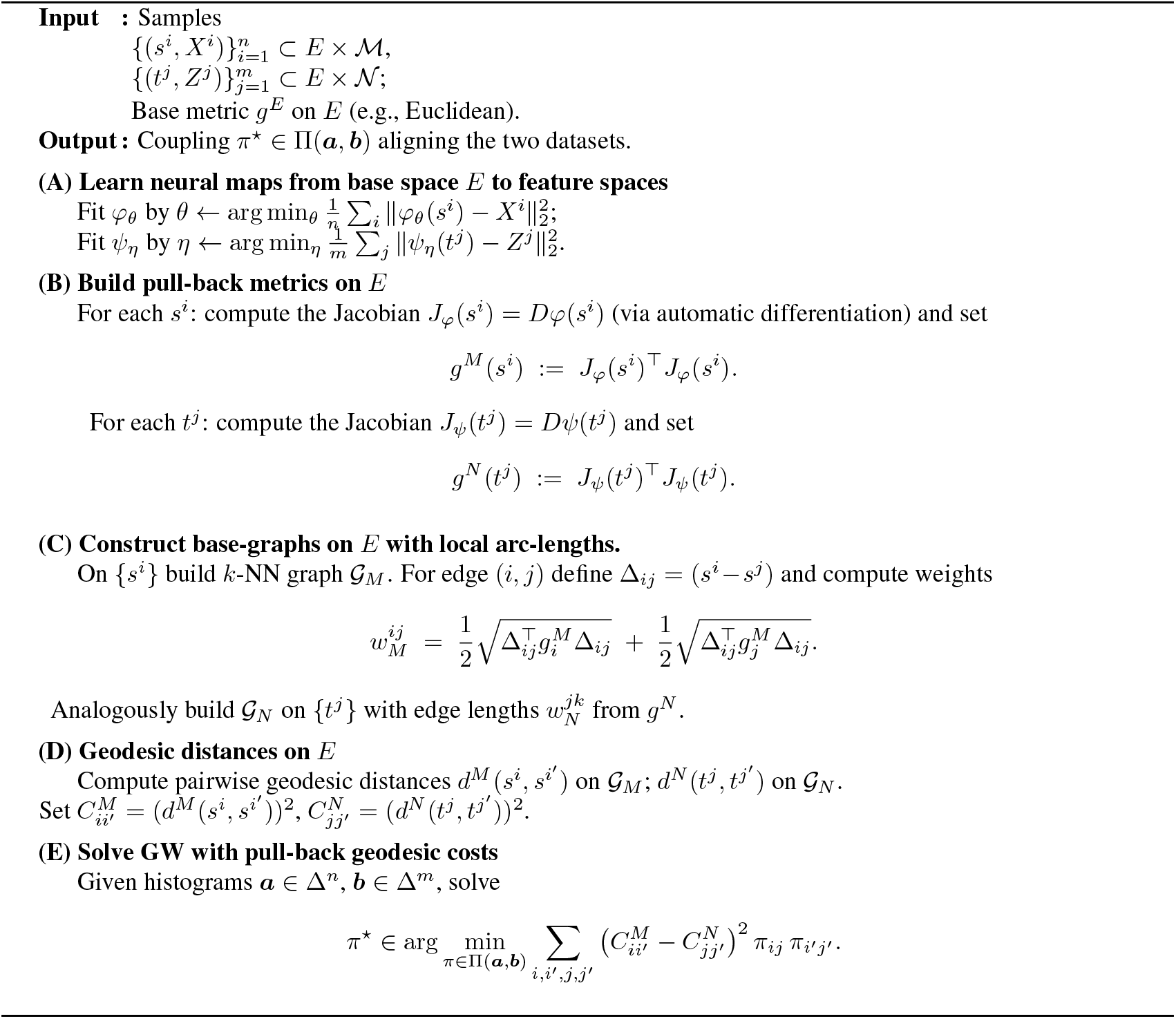

### C. Experimental Details

#### C.1 Datasets, Availability, and Pre-processing

In Section 3, we have evaluations of: (1) a spatiotemporal mouse transcriptomics Stereo-Seq dataset (5) (Section 3.1), an evaluation of a Visium 10x Genomics colorectal cancer (CRC) dataset and a Xenium dataset at the same slide (37) (Section 3.2), (3) an alignment of a MALDI-MSI metabolomics dataset and Visium transcriptomics dataset of a human striatum brain section (56) (Section 3.4), and (4) and alignment of an AFADESI-MSI metabolomics and Visium transcriptomics dataset of human clear cell renal carcinoma (ccRCC) (53) (Section 3.3). All datasets are publicly accessible, and we publish the code for loading and processing each.

##### Stereo-Seq Spatiotemporal Transcriptomics

For the dataset of (5) we use `scanpy` to load AnnData files for each day of mouse embryonic development from E9.5 to E13.5

- `E9.5_E1S1.MOSTA.h5ad`
- `E10.5_E1S1.MOSTA.h5ad`
- `E11.5_E1S1.MOSTA.h5ad`
- `E12.5_E1S1.MOSTA.h5ad`
- `E13.5_E1S1.MOSTA.h5ad`

We construct paired datasets by taking the intersection of common genes across the timepoints, and perform a joint PCA across the time pairs. To do this, we concatenate each consecutive timepoint pair and normalize the joint AnnData with `sc.pp.normalize_total` and add pseudo-counts with `sc.pp.log1p` for feature-sparsity. We then perform PCA on the joint AnnData with 30 components. As the PCA is performed jointly in this experiment, no additional CCA step is required for feature-alignment. Owing to the size of the slices, we randomly downsample each to 10k points with a random seed of `rng=`42.

##### Visium-Xenium Alignment

For the public dataset of (37) we use the colorectal cancer (CRC) dataset. This consists of two multi-modal sections from the same donor (Sample P2 CRC) (37). This dataset includes a Visium CytAssist v2 section and a Xenium spatial transcriptomics section profiling the same tissue. We use this dataset, as it exhibits significant overlap (≈ 90%, (61)), making it an appropriate candidate for alignment. As the Visium technology offers sequencing of the transcriptomics (≈ 18, 000 genes), and Xenium offers FISH-based imagining (422 genes), we treat the datasets as distinct modalities. As a result, we perform independent PCA on each dataset (as opposed to a joint PCA) with 100 components. We then perform a CCA, as outlined in Section C.6, with 40 components to find the most mutually correlated components within the independent PCA sets.

##### Transcriptomics-Metabolomics of Human Striatum

We download the Visium and MALDI-MSI datasets for the slide `V11T17-102_A1` from (56), corresponding to a post-mortem slice of human striatum in with Parkinson’s diseases. The data deposition of (56) is available at Mendeley Data through the link https://data.mendeley.com/datasets/w7nw4km7xd/1. We select this slide in particular, as it is the only dataset with published annotations for (dopamine-annotated) Visium neurons which we use as a ground-truth in our benchmarking. These are contained in `dopamine.csv` in the directory `V11T17-102/V11T17-102_A1/output_data/V11T17-102_A1_RNA/outs/dopamine.csv`. We use the label `dopamine_Cd` as the ground-truth dopamine predictor in the caudate-nucleus (Cd), and use the labels `not_dopamine_Cd` and `CI` as the negative labels. We pre-process the metabolomics dataset with filtering for the top spatially-varying metabolites. We first normalize the metabolomics data with scanpy as `sc.pp.normalize_total`, and then use squidpy to compute the spatially variable metabolites with `sq.gr.spatial_neighbors` and `sq.gr.spatial_autocorr`, using `mode=“moran”` to employ the Moran’s I statistic. We select the top 20 metabolites in this step, and use this pre-processed data for all methods. For `MGW` we do not perform any additional processing of the raw metabolomics data beyond filtering. We use the CCA step (Section C.6) to align the metabolomics and transcriptomics (PCA) features to 3 CC components in a joint-space suitable for `MGW` and use these features for alignment.

##### Transcriptomics-Metabolomics of Renal Cancer

To benchmark against the multimodal model `SpatialMeta`, we employed the clear cell renal cell carcinoma (ccRCC) dataset from its original publication (53). The processed spatial transcriptomics (ST) and spatial metabolomics (SM) data for the `Y7_T` tissue section were obtained from the Zenodo repository (accession code 14986870), as `Y7_T` was the only slice for which both raw ST and SM data were publicly available. The corresponding ST-SM alignment generated by `SpatialMeta` was also downloaded from the same source.

For our method (`MGW`) and other optimal transport-based baselines, we followed `SpatialMeta`’s preprocessing workflow provided in their official GitHub repository (Vertical_and_Horizontal_Integration_for_ST_SM.ipynb). To ensure a fair comparison, we adopted the same variational autoencoder (VAE) architecture as `SpatialMeta` across all methods and trained each model for 100 epochs to obtain the low-dimensional embeddings. We repeat the evaluation of the entire across three random seeds to ensure reproducibility.

Model evaluation was performed using `SpatialMeta`’s official benchmarking pipeline. In particular, this was done via the function `run_modality_benchmark` which is implemented in `multi_benchmark_function.py` in the `benchmark/` folder. Since the manual pathological annotations referenced in the original publication were not included in the released dataset, metrics dependent on these annotations were excluded from our evaluation, and the remaining quantitative metrics were computed consistently across all models.

#### C.2 Baselines

We benchmark `MGW` against the methods `moscot` (26), `SCOT` (11), `SCOTv2` (10), SpatialMeta (53), and PASTE2 (30), in addition to two baselines: Gromov-Wasserstein optimal transport on (1) features and (2) spatial coordinates. The feature input is standardized for all methods to the PCA components computed by `mgw.mgw_preprocess`. We run `moscot moscot.problems.TranslationProblem` with `max_iter=5_000` and `tol=1e-7`. Following the tutorial, available at https://moscot.readthedocs.io/en/latest/notebooks/tutorials/600_tutorial_translation.html, we use the default `alpha = 1` as one must ignore the `joint_attr` for multiomics alignment. For `SCOT` we following the procedure outlined in the tutorial https://rsinghlab.github.io/SCOT/tutorial/. We set `k=50` for the nearest neighbor graph, `epsilon=0.005` for the entropic regularization, and set `normalize=True`. We use the same parameters for `SCOTv2`, with the additional parameter of `rho=0.1`. We run `PASTE2` with `s=0.7` and replace the `adata.X` coordinate with spatial coordinates, as multiomic alignment lacks joint distances across features. Lastly, for the Gromov-Wasserstein baselines, we use the package `ott-jax` (9) with feature and spatial costs constructed with squared-Euclidean distance as 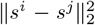 and 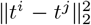 (spatial), as well as 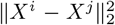 and 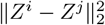 (feature in spaces ℳ and 𝒩). We use the parameters `inner_maxit=3000`, `outer_maxit=3000`, `inner_tol=1e-8`, `outer_tol=1e-8`, `epsilon=1e-4` (Section 3.1), exactly paralleling the settings of `ott-jax` used for `MGW` but with a different cost. For the comparison to `SpatialMeta`, we downloaded their aligned `adata_joint_Y7_T_raw.h5ad` for `Y7_T` slice from their published Zenodo source https://zenodo.org/records/14986870. Then we trained the `MGW`, `SpatialMeta` and other models on `ConditionalVAESTSM` from `SpatialMeta`’s Github for 100 epochs to retrieve the embedding for downstream analysis.

#### C.3 Architecture and Parameters

To train our neural fields φ: ℝ^2^ → ℳ and *ψ* : ℝ^2^ → 𝒩, we use multi-layer perceptrons (MLPs). For *ψ*, φ to be smooth sub-manifolds with sufficiently smooth Jacobians we require smooth *C*^∞^ activations: in particular, we exclude activation functions like ReLU and use the SiLU or SWISH function (41; 35). We use an MLP with widths (128, 256, 256, 128), learning rate *η* = 1 × 10^−3^, weight-decay of *w*_decay_ = 1 × 10^−4^, and EMA (exponential moving average) decay of EMA_decay_ = 0.995. We train the network for a total of `niter=20_000` training iterations.

#### C.4 Computing metric tensor-field and geodesics

We use `vmap(jacrev(phi))(X)` to compute Jacobians at each point, and compute the pull-back metric tensor field using an Einstein summation of the form `torch.einsum(‘nfd,nfe->nde’, J, J)`. We normalize this tensor-field across space with the mean-eigenvalue of *g* as a proxy, and add a small stabilization of the form 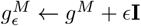. As *g*^*M*^ is scaled ≈ 1 we typically choose *ϵ* = 10^−2^ to be much smaller, but sufficiently large to avoid instabilities for under-determined *g*.

To practically realize geodesics, we compute a *K*-nearest neighbor graph. On spatial positions { *s*^*i*^ } we build *k*-NN graph 𝒢_*M*_. For edge (*i, j*) define Δ_*ij*_ = (*s*^*i*^ − *s*^*j*^). Then, the edge-weights in 𝒢_*M*_ are given by the arc-lengths in the Riemannian metric

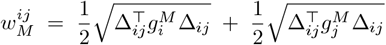

With *g*^*M*^ defined from the neural field at each point *g*^*M*^ (*s*^*i*^)(·,·) and *g*^*M*^ (*s*^*j*^)(·,·). From this graph, geodesic distances are computed with Dykstra’s algorithm from `scipy.sparse.csgraph.shortest_path`.

#### C.5 Details on the Optimal Transport

For the optimal transport, we use the Gromov-Wasserstein solver of `ott-jax` (9). This relies on using our instantiated geodesic costs as

- `geometry.Geometry(cost_matrix=CM)`
- `geometry.Geometry(cost_matrix=CN)`and calling `quadratic_problem.QuadraticProblem` on these objects. We use parameters of `inner_maxit=3000`, `outer_maxit=3000`, `inner_tol=1e-8`, `outer_tol=1e-8`, and `epsilon=1e-4`. We offer the unbalanced setting with *τ*`_a` and *τ*`_b` as options, but set both to 1 (Balanced) for all experiments.

#### C.6 Pre-processing for the Alignment-Informative Feature Subset

The feature spaces of distinct modalities need not align nor exhibit meaningful joint structure. There may be “marginal” sub-spaces of each feature which may be independent of the other modality, and there may be “joint” sub-spaces which exhibit high-correlation across the modalities. Thus, a natural pre-processing is to filter for such joint sub-spaces.

We propose a step based on *canonical correlation analysis* (CCA) (20), which identifies linear combinations of two datasets which are maximally correlated. Formally, given random vectors *X* ∈ ℝ^*n*^, *Y* ∈ ℝ^*m*^ CCA computes a sequence of vectors 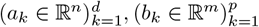 that maximize the correlation of their projections 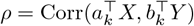. To isolate the joint sub-spaces, we propose an initial feature-refinement step comprising two components.

*First*, compute a coarse alignment 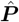 between the two datasets **X** and **Z**: an approximate correspondence is sufficient to identify features with strong cross-modal correspondence. *Second*, one takes a barycentric projection of the second dataset onto the first 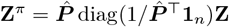. This ensures a 1-1 alignment of the points, which is a requirement of CCA. Following projection the canonical components are obtained by solving arg 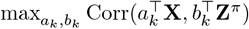 for *k* ∈ [*d*]. This yields canonically loading matrices **U**_*a*_, **U**_*b*_ with columns representing the maximally correlated directions of the two modalities. We then project the features in both modalities to their maximally-correlated components prior to `MGW` alignment. For a common modality, we perform a joint PCA across datasets in place of CCA.

### D Supplementary Figures and Tables

**Figure S7:**
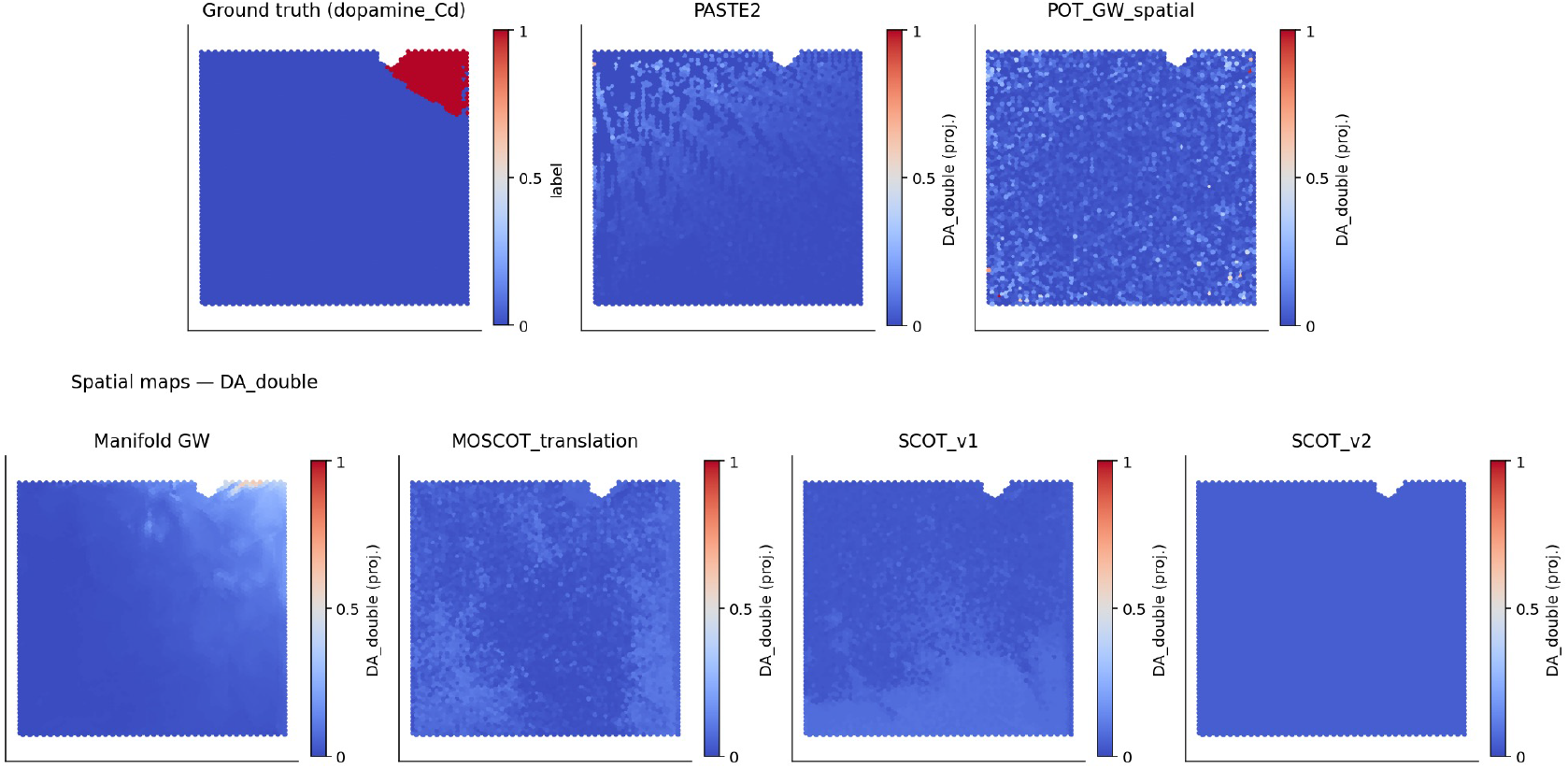
Raw, unscaled DA Double (doubly-derivatized dopamine, *m/z* = 674.28) following barycentric projection of metabolite intensities onto Visium Slide of (56) across couplings ***P*** returned by various methods.

**Figure S8:**
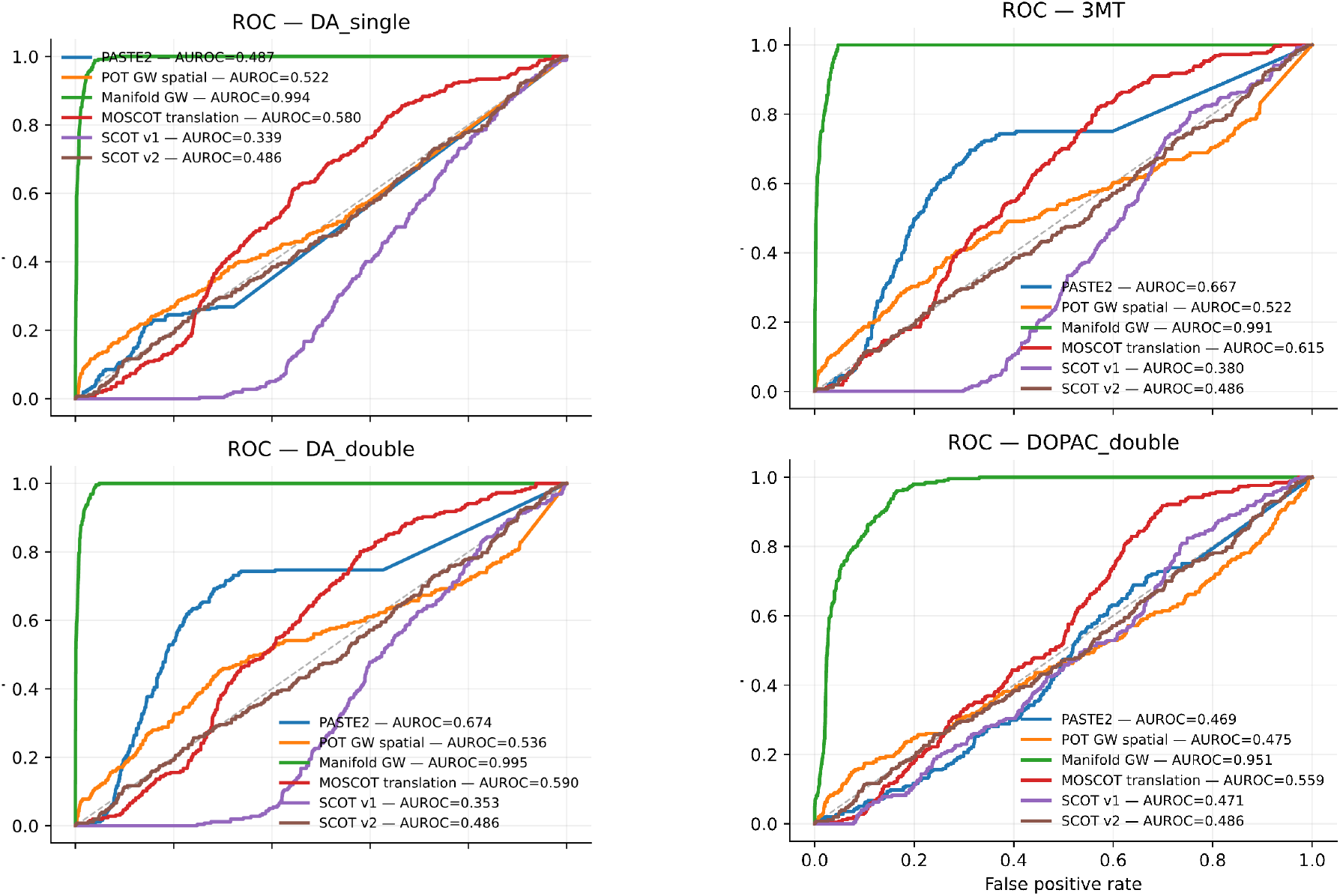
AUROC curves for the 4 dopamine metabolites DA Single (singly-derivatized *m/z* = 421.19), DA double (doubly-derivatized *m/z* = 674.28), DOPAC double (dopamine-breakdown product *m/z* = 698.24), and 3MT (dopamine-breakdown product 3-Methoxytyramine, *m/z* = 435.21) in the MALDI-MSI to Visium Transfer Task.

**Figure S9:**
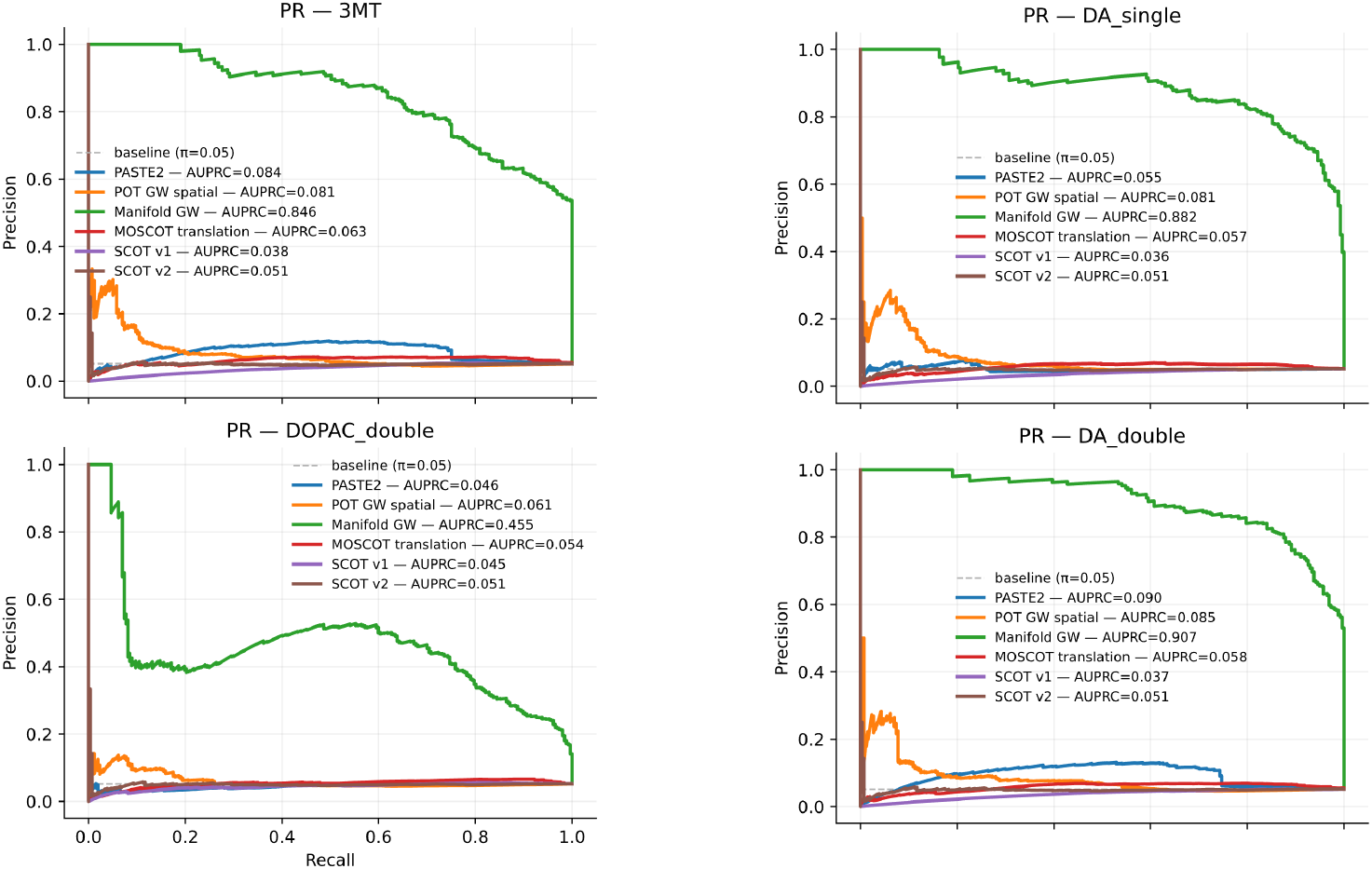
AUPRC curves for the 4 dopamine metabolites DA Single (singly-derivatized *m/z* = 421.19), DA double (doubly-derivatized *m/z* = 674.28), DOPAC double (dopamine-breakdown product *m/z* = 698.24), and 3MT (dopamine-breakdown product 3-Methoxytyramine, *m/z* = 435.21) in the MALDI-MSI to Visium Transfer Task.

**Figure S10:**
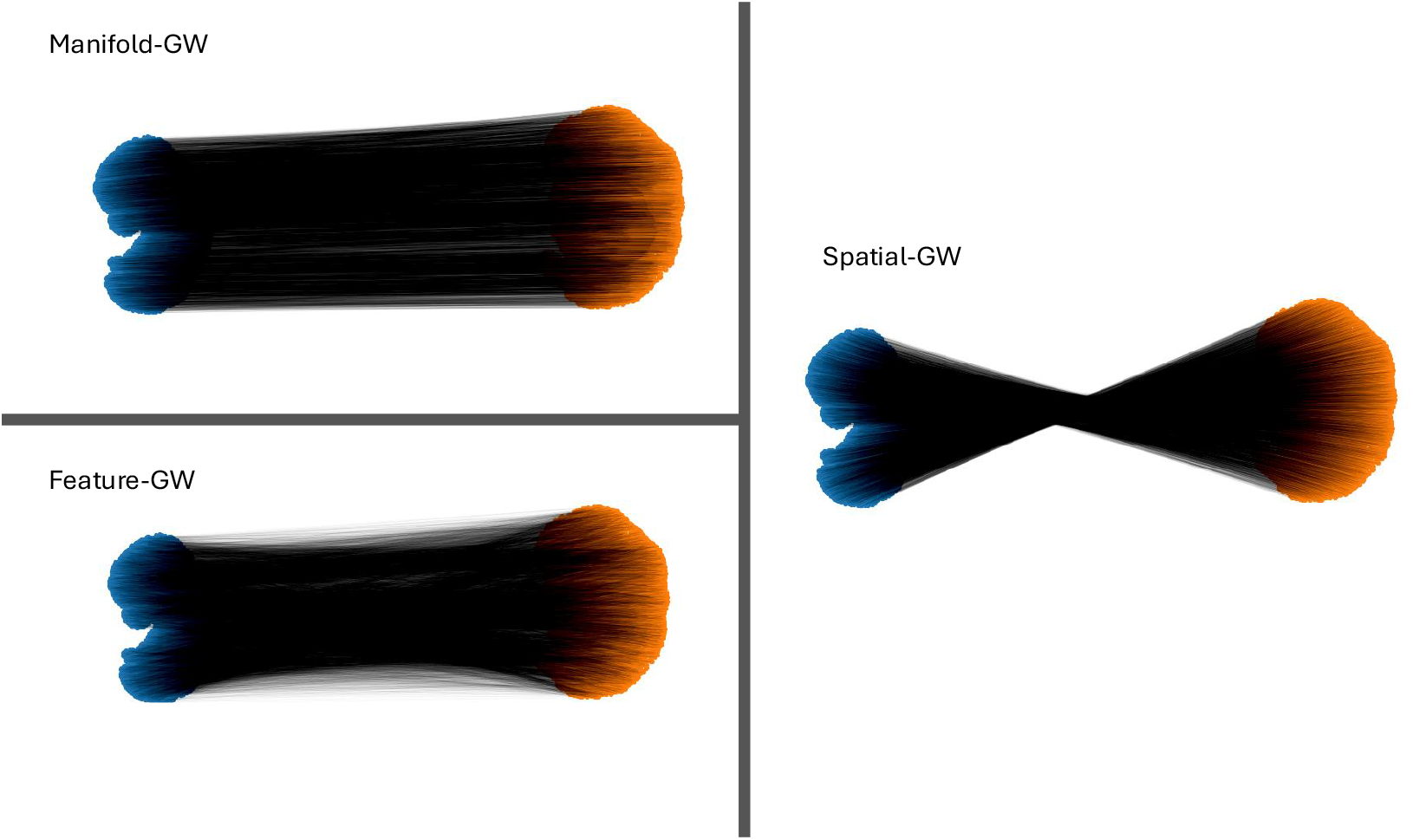
Visualization of alignments computed by Manifold-GW, as well as Gromov-Wasserstein baselines Spatial-only GW and Feature-only GW. Alignments shown for Stereo-seq mouse-embryo timepairs E12.5-13.5 from (5).

**Figure S11:**
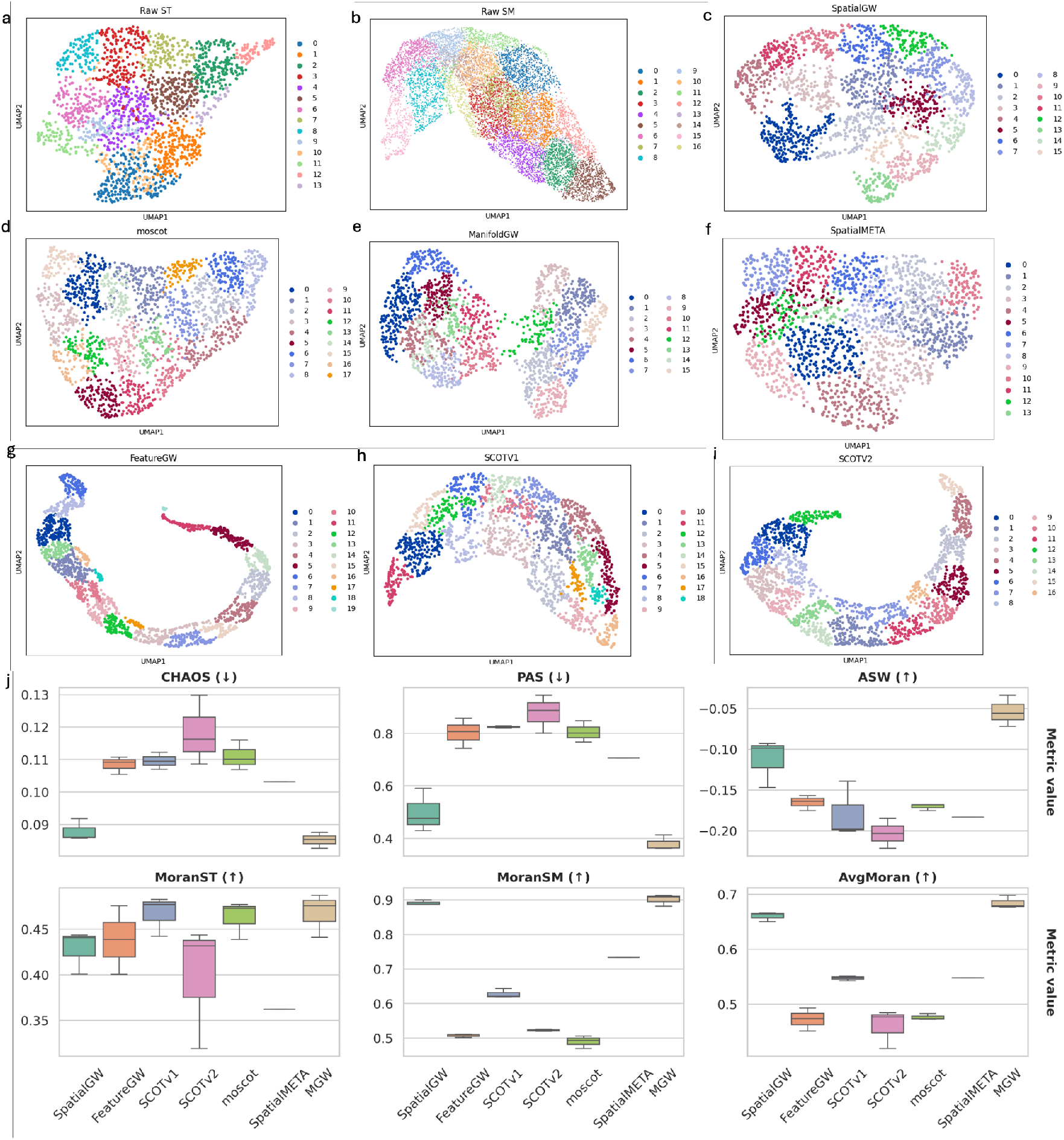
**(a-i)** The UMAP visualization of the aligned embeddings of *Y* _*T* slice from ccRCC dataset of all models.**(j)** The boxplot of all six metrics across three seeds as in table 2. The `SpatialMeta`’s value is consistent across three seeds, because it is not an optimal transport based model and there is no randomness coming from the approximation of coupling (53). The up or down arrow next to the metric stands for if a higher or lower value means better performance.

## Footnotes

1 The cost may depend on the coupling, as in (11).

2 Or, equivalently, a permutation *σ*: (*s*^*i*^, *X*^*i*^) → (*t*^*σ*(*i*)^, *Z*^*σ*(*i*)^)

